# Cholinergic basal forebrain degeneration due to obstructive sleep apnoea increases Alzheimer’s pathology in mice

**DOI:** 10.1101/2020.03.12.989848

**Authors:** Lei Qian, Leda Kasas, Michael R Milne, Oliver Rawashdeh, Nicola Marks, Aanchal Sharma, Mark C Bellingham, Elizabeth J Coulson

**Affiliations:** Queensland Brain Institute, The University of Queensland, Brisbane Qld. 4072, Australia; Clem Jones Centre for Ageing Dementia Research, The University of Queensland, Brisbane Qld. 4072, Australia; Faculty of Medicine, School of Biomedical Sciences, The University of Queensland, Brisbane Qld. 4072, Australia

**Keywords:** obstructive sleep apnoea, intermittent hypoxia, basal forebrain, cholinergic neuron, Alzheimer’s disease, hypoxia-inducible factor 1alpha

## Abstract

Epidemiological studies indicate that obstructive sleep apnoea is a strong risk factor for the development of Alzheimer’s disease but the mechanisms of the risk remain unclear. We developed a method of modelling obstructive sleep apnoea in mice that replicates key features of human obstructive sleep apnoea: altered breathing during sleep, sleep disruption, moderate intermittent hypoxemia and cognitive impairment. When we induced obstructive sleep apnoea in a familial Alzheimer’s disease model, the mice displayed exacerbation of cognitive impairment and pathological features of Alzheimer’s disease, including increased levels of amyloid-beta and inflammatory markers, as well as selective degeneration of cholinergic basal forebrain neurons. These pathological features were not induced by chronic hypoxia or sleep disruption alone. Our results also revealed that the neurodegeneration was mediated by the oxygen-sensitive p75 neurotrophin receptor and hypoxia inducible factor 1 alpha activity. Furthermore, restoring blood oxygen levels during sleep to prevent intermittent hypoxia prevented the pathological changes induced by the OSA. These findings provide a signalling mechanism by which obstructive sleep apnoea induces cholinergic basal forebrain degeneration and could thereby increase the risk of developing Alzheimer’s disease, as well as providing a rationale for testing a range of possible prophylactic treatment options for people with obstructive sleep apnoea and hypoxia including increased compliance of continuous positive airway pressure therapy.

## Introduction

The causes of idiopathic dementia, of which Alzheimer’s disease (AD) is the largest subgroup, are not clear. The leading theory of AD is based on familial forms of the condition in which the cleavage of amyloid precursor protein (APP) is altered, favouring production of the neurotoxic peptide amyloid-beta (Aβ) (Hardy and Selkoe, 2002). However, the physiological triggers that lead to change to APP metabolism, or that result in Aβ accumulation, are poorly defined (Hardy and Selkoe, 2002). Furthermore, Aβ accumulation alone is not sufficient to induce cognitive decline in the elderly. Elucidating the aetiology of idiopathic AD is therefore crucial to providing efficacious early intervention and/or treatment for the majority of people who develop AD.

Obstructive sleep apnoea (OSA) is a strong epidemiological risk factor for the development of dementia (Zhu and Zhao, 2018). It affects more than 50% of the elderly adult population (Punjabi, 2008) and occurs due to the collapse of the upper airways, particularly during rapid eye movement (REM) sleep; this impedes airflow and requires a brief period of wakefulness in order to re-activate airway tone (Yaffe et al., 2014; Daulatzai, 2015). OSA has been associated with both an earlier age of AD onset, more rapid cognitive decline (Ancoli-Israel et al., 2008; Osorio et al., 2011; Yaffe et al., 2014) and an increased Aβ burden as measured by PET imaging (Yun et al., 2017; Elias et al., 2018). However, OSA is not widely recognized as one of the lifestyle factors presenting increased risk of developing AD (Livingston et al., 2017).

Although the reasons for the epidemiological association between OSA and AD are unclear, there are a number of non-mutually exclusive possibilities. For example, longitudinal studies suggest that the hypoxia which results from OSA causes neurodegeneration, thereby leading to cognitive decline (Yaffe et al., 2011; Daulatzai, 2015). Similarly, exposure of mice to intermittent hypoxia can exacerbate the accumulation of Aβ (Shiota et al., 2013) and cause neurodegeneration (Row et al., 2007). However, a more recent hypothesis is that disrupted sleep slows the clearance of Aβ from the brain interstitial space through the glymphatic system (Mander et al., 2016). Finally, sleep, in particular REM sleep, is considered fundamental for learning and memory consolidation, and poor sleep quality leads to failed memory consolidation (Wang et al., 2011). Therefore OSA-induced sleep disruption could contribute to cognitive impairment and eventually dementia.

The reasons for the increased risk that OSA creates are difficult to determine in humans, even in longitudinal studies, due to the high comorbidity rates of other risk factors such as diabetes and cardiovascular disease in people suffering from OSA, most of whom are also overweight or obese (Livingston et al., 2017). In order to understand the mechanisms linking AD with OSA we developed a method of modelling OSA in mice in the absence of comorbidities. We then studied the possible underlying mechanisms by which OSA affects the aetiology of key hallmarks of AD.

## Materials and Methods

### Animals

C57BL/6, APP/PS1(JAX - 34832), ChAT-cre x p75_fl/fl_ (Boskovic et al., 2014) or TrkA-cre x p75_fl/fl_ (Sanchez-Ortiz et al., 2012; Boskovic et al., 2019) mice were maintained on a 12h light/dark cycle with *ad libitum* access to food and water. Mice of either sex were used, unless otherwise indicated. Littermates of the same sex were randomly assigned to experimental groups and used at 8-12 weeks or 8 months (for APP/PS1) of age unless stated otherwise. All procedures were approved by the University of Queensland Animal Ethics Committee and conducted in accordance with the Australian Code of Practice for the Care and Use of Animals for Scientific Purposes (8th edition, 2013).

### Surgery

Standard surgical procedures were followed for stereotaxic injection. To induce cholinergic mesopontine tegmentum (cMPT) lesions, bilateral injections of urotensin II-saporin (UII-SAP; 0.07μg/μL per site – unless stated otherwise; generous gift from Advanced Targeting Systems) in the laterodorsal tegmental nucleus (A-P, −3.0 mm; M-L, ±0.5 mm; D-V, −4.0 mm from Bregma, with the angle 34° backward) were made using a calibrated glass micropipette through a Picospritzer ® II (Parker Hannifin), with the same molar mass of blank-saporin (Blank-SAP) or IgG-saporin (IgG-SAP) as control.

### Phenotypic analysis

Behavioural experiments were always performed in the light cycle. Some cohorts of mice were subject to multiple behavioural tests.

#### Sleep deprivation

To induce sleep deprivation, up to 8 mice were placed in a water-filled cage (55cm x 25cm) that contained 9 cylinders (3cm diameter) with 1 cm above the water surface (Sup Fig 7). The sleep deprivation cages and control cages, which also contained cylinders but no water inside, were held within a climate chamber on a 12:12 h light:dark cycle at 27 °C and 25% relative humidity. Mice were placed in the sleep deprivation cages for 20 h a day and in their home cages (4 to a cage) also within the cabinet, for 4 h per day during their light cycle. Mice were monitored by video and weighed daily.

#### Whole body plethymography

Whole body plethysmography has been used to record respiration during sleep and wake cycles in unrestrained, freely moving mice (Hernandez et al., 2012). The mice were placed in a plethysmograph chamber (Buxco FinePointe Series WBP, DSI) continuously filled with fresh air at room temperature. This approach provides an indirect measure of tidal volume, which is directly proportional to the cyclic chamber pressure signal produced during respiration in a sealed chamber. The mice were first allowed 30 min to acclimatize to the chamber, after which the recording session preceded for 3h.

#### Arterial oxygen saturation

The arterial oxygen saturation was recorded on unrestrained awake mice either in room air or in the environmental chamber using the MouseOx Plus oximeter (Starr Life Sciences) in accordance with the manufacturer’s instructions. Data were collected for at least 30 min and only used when no error code was given. Oxygen saturation levels were then calculated as an average per mouse, and per experimental condition.

#### Open field

The open field test was performed in a square white Plexiglas box (30cm x 30cm) for 30 minutes. The chamber was divided into a central field (centre, 15cm x 15cm) and an outer field (border). The mouse movement was recorded using a video camera, and analysed using the EthoVision XT video tracking system (Noldus Information Technology). Total track length was calculated from the centre of the animal’s body as the reference point.

#### Y maze

The Y-maze was composed of three equally spaced arms (120° apart, 35 cm long and 10cm wide) made of Perspex. During training the mice were placed in one of the arms (the start arm) with access to only one other arm for a period of 15 min. One day after the training, mice were allowed to explore in the Y-maze for 10 min with access to all the three arms. Animals were tracked using EthoVision XT software and the total time and the percent time spent in each arm were analysed.

#### Novel object recognition

The task procedure consisted of three phases: habituation, familiarization, and the test phase. In the habituation phase, the mice were allowed to freely explore the open-field arena (40 cm x 40 cm) for 5 min. During the familiarization phase, they were placed in the arena containing two identical sample objects, for 5 min, where both objects were located in opposite and symmetrical corners of the arena. After a 1 h retention interval, the mice were returned to the arena with two objects, one identical to the sample and the other novel, and were allowed to explore again for 5 min. Animals were tracked using EthoVision XT software and the percent time spent exploring the objects were analysed.

#### Active place avoidance

The apparatus (Bio-Signal Group) consisted of an elevated arena with a grid floor fenced with a transparent circular boundary, located in a room with visual cues on the walls (Cimadevilla et al., 2000; Wesierska et al., 2005). The arena rotated clockwise (1 rpm) and an electric shock could be delivered through the grid floor. On day after the habituation, mice underwent daily training sessions for 4 or 5 days, lasting for 10 min per day in which the mouse was trained to avoid a 60° shock zone, where entrance led to the delivery of a brief foot shock (500 ms, 0.5 mA). To determine their memory of the shock zone, 24 h after the last training session, the mice were allowed to explore the arena for 10 min without shocks being applied. The position of the animal in the arena was tracked using an overhead camera linked to Tracker software (Bio-Signal Group).

#### Passive place avoidance

The passive place avoidance behavioural paradigm was used to test basal forebrain-dependent idiothetic navigation (Cimadevilla et al., 2000; Hamlin et al., 2013). This navigation task used the same apparatus as the active place avoidance task. However, all external cues were eliminated to minimize allothetic navigation, the circular boundary was of a grey opaque and the arena remained stationary throughout the experiment. The mice went through a 5-min habituation session and then a 10-min training phase with 1 hr interval between them. Twenty-four hr after the training session, the mice were allowed to explore the arena for 5 min without shocks being applied zone (Hamlin et al., 2013).

#### Morris water maze

Mice were trained to escape the maze by finding the platform submerged 1.5 cm beneath the surface of the water and invisible to the mice while swimming. Three trials were performed per day, for 60 s each with 30 min between trials. During each trial, mice were placed into the tank at one of three designated start points in a pseudorandom order. If the mice failed to find the platform within 60 s, they were manually guided to the platform and allowed to remain there for at least 20 s. Mice were trained for 5 days as needed to reach the training criterion of 15 s (escape latency). The probe trial occurred 24 h after the last training session and consisted of a 60 s trial without the platform. Performance was tracking with the EthoVision XT video tracking system.

#### Sleep analyses using the PhenoMaster

The locomotor activity and sleep status of the mice were assessed using metabolic cages PhenoMaster (TSE Systems). The mice were acclimatized in the PhenoMaster for 4 days before the start of the recordings. The mice were continuously recorded for at least 3 days and the activity of each mouse was monitored using infrared motion sensors. For the assessment of effects of individual light pulses to induce sleep in the dark, three light treatments of 100, 50, or 30 lux were applied during the dark phase of the light cycle for 3 h at the day 4, 7, or 10, respectively. Following the sleep deprivation experiment, the mice were individually housed and were continuously recorded for 48 h after the completion of sleep deprivation treatment without habituation.

### 2ME2 treatments

Two weeks post-surgery, the cMPT-lesioned mice were treated with either a HIF1α inhibitor 2-methoxyestradiol (2ME2, Sigma-Aldrich, M6383), or vehicle control (saline with 0.5% DMSO). The 2ME2 was dissolved in saline at a concentration of 0.5 mg/mL with final 0.5% DMSO. Mice received daily i.p. injections of either 2ME2 (15 mg/kg body weight) or vehicle for 3 or 4 weeks. Mice were weighed every third day to detect any body weight changes and possible overt toxicity of long-term 2ME2 administration (no changes were observed).

### Immunohistochemistry

For immunofluorescence labelling, free-floating sections were probed using the following primary antibodies: goat anti-ChAT (1:400, AB144P, Millipore), mouse anti-parvalbumin (1:1000, MAB1572, Millipore), rabbit anti-calbindin (1:2000, CB38, Swant), mouse anti-Aβ (6E10, 1:500, Sig-39320, Convance), rabbit anti-GFAP (1:500, Z0334, Dako) and rat anti-CD68 (FA-11, 1:500, MCA1957, AbD Serotec), rabbit ant-HIF1α (1:200, NB100-479, Novus Biologicals) followed by the appropriate secondary antibody (1:1000, Life Technologies) or incubated with thioflavin S (0.1% in water, T1892, Sigma-Aldrich). Sections were mounted onto slides and coverslipped using fluorescence mounting medium (Dako).

For DAB (3,3’-diaminobenzidine) staining, the biotinylated donkey secondary antibodies (1:1000, Jackson ImmunoResearch Laboratories) and ABC reagent (Vector Laboratories) were applied following the primary antibody incubation and followed by the nickel-intensified DAB reaction. Brain slices were mounted on slides and coverslipped with DPX mounting medium (Sigma-Aldrich).

#### ELISA methods

For biochemical analyses, mice were perfused with PBS then the cortex and hippocampus were dissected, weighed, and snapfrozen with liquid nitrogen. The protein was extracted by adding ice-cold 10 v/w RIPA buffer (250 mM NaCl, 1% NP-40, 0.5% sodium deoxycholate, 0.1% SDS, 50 mM Tris HCl, pH 8.0, containing Roche cOmplete™ protease inhibitor cocktail and PhosSTOP phosphatase inhibitor cocktail and 1mM AEBSF) to the tissue before homogenization using Bullet Blender Storm 24 (Next Advance).

The level of Aβ in the tissue was assessed by ELISA as per the manufacturer’s instructions (KHB3442, Invitrogen). The resulting measurements were normalized for tissue weight.

### Statistical analysis

Statistical analysis was performed using GraphPad Prism 7 software, accounting for appropriate distribution and variance to ensure proper statistical parameters were applied. Values are expressed as the mean ± s.e.m with significance determined at p < 0.05. Data from different cohorts of mice that had undergone the same procedures were combined for analyses. Mice without a cMPT lesion were excluded from analysis.

### Image analysis and histological quantification

Imaging was obtained using either an upright fluorescence microscope (Axio Imager, Zeiss) and VSlide Scanner (Metafer, Metasystems), or a Diskovery spinning disk confocal microscope (Andor, Oxford Instruments). All measurements and analyses were performed using Imaris 9.2.1 software (Bitplane) or Image J.

### Data availability

The data that support the findings of this study are available from the corresponding author upon request.

## Results

### Urotensin 2-saporin toxin specifically lesions cholinergic mesopontine neurons, resulting in the first animal model of OSA

Mesopontine tegmentum (MPT) neurons innervate brain regions important in REM sleep, such as thalamocortical regions and the basal forebrain (Woolf and Butcher, 1986), and are strongly implicated in initiating and maintaining REM sleep (Zhang et al., 2005; Van Dort et al.). Cholinergic MPT (cMPT) neurons also project to the hypoglossal motor nucleus, which controls the tongue muscles, as well as to other brainstem areas (Woolf and Butcher, 1986). Respiratory activity in upper airway muscle motor neurons is driven by the brain stem respiratory central pattern generator (Bellingham and Berger, 1996), and the tongue muscles are important upper airway dilator muscles during inspiration (Mizuno et al., 2004). We reasoned that cMPT lesioning would result in upper airway collapse, particularly during REM sleep when acetylcholine activity is highest (Irmak and de Lecea, 2014), providing a naturalistic mouse model of OSA.

cMPT neurons selectively express the urotensin 2 receptor, and respond to its ligand, urotensin-2 peptide (Clark et al., 2001; Clark et al., 2005; Jegou et al., 2006). To selectively ablate cMPT neurons we used a ribosomal inactivating saporin-conjugated urotensin-2 peptide (UII-SAP) targeting the lateral dorsal tegmentum (LDT) region rather than the pedunculopontine nucleus (PPN) of the MPT, which is known to mediate locomotion or reward (Xiao et al., 2016). Direct injection of the UII-SAP toxin into the brain stem induced a unilateral specific loss of cholinergic neurons within the MPT after 2 weeks (**Fig 1**; the dose response of UII-SAP is shown in **Supplementary Fig 1A**). Compared to a control injection of rabbit-IgG-saporin (IgG-SAP; not shown) or blank-saporin (Blank-SAP) which is conjugated with a non-targeted peptide consisting of similar, comparable materials, the number of choline acetyltransferase (ChAT)-positive neurons in the LDT was significantly reduced in UII-SAP injected mice (**Fig 1A,B**), with a smaller but significant loss of ChAT-positive neurons also being detected in the PPN. A similar loss of cLDT neurons was observed when UII-SAP was injected via the intracerebral ventricle (**Fig 1C**), indicating that the reduced number of ChAT-positive neurons was not due to damage at the injection site. However, there was no loss of calbindin-positive GABAergic neurons in the MPT (**Fig 1D**) or of cholinergic hypoglossus motor neurons (**Fig 1E**). cMPT-lesioned mice displayed no significant change in body weight (not shown), the distance travelled (**Fig 1F**) or exploration (**Sup. Fig 1B**) in an open field test, or in rotarod performance (**Fig 1G**), relative to control-lesioned mice.

**Figure 1.**
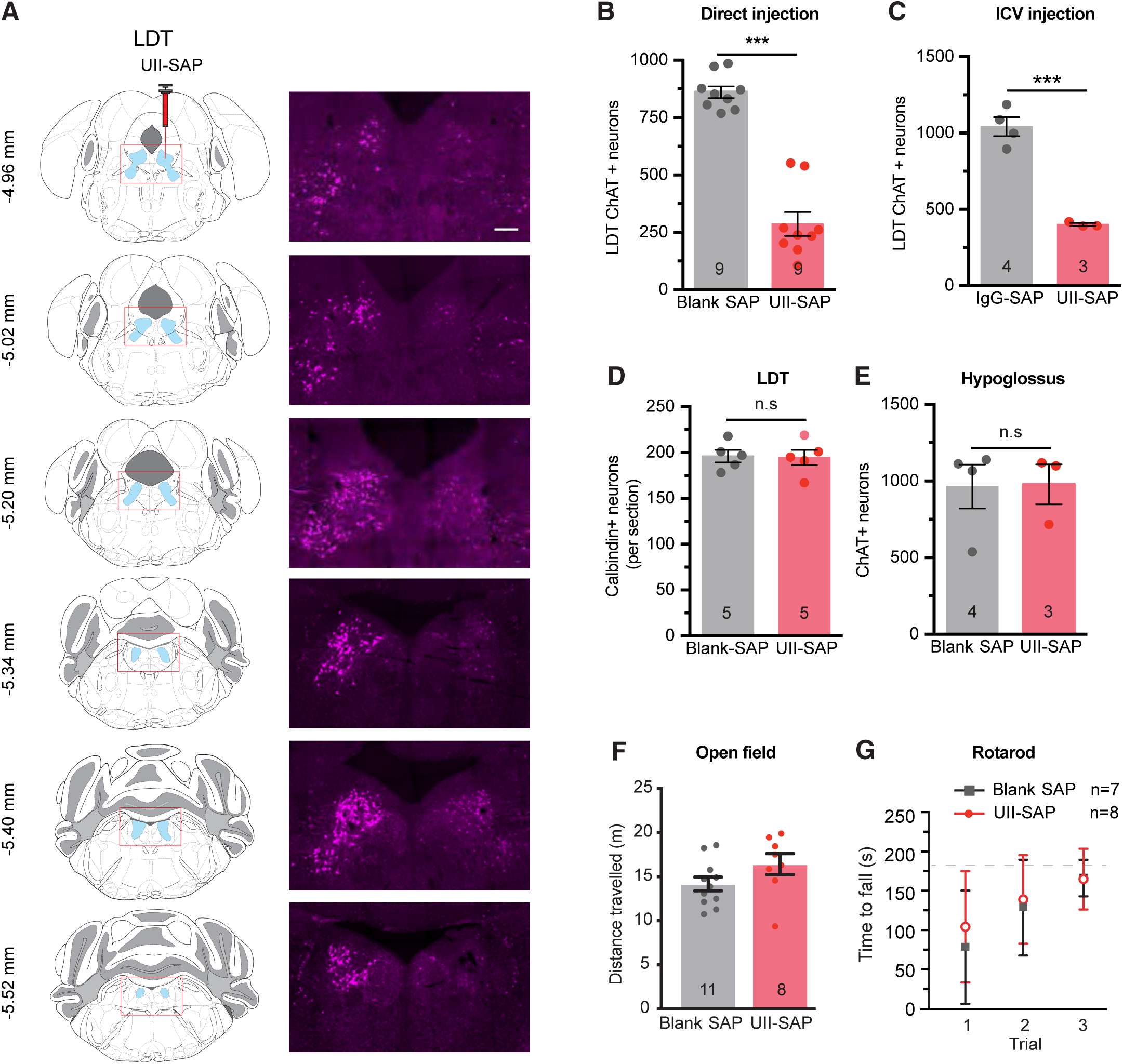
Urotensin II-saporin induces specific lesions of cholinergic neurons at mesopontine tegmentum. **(A)** Representative diagrams and photomicrographs of coronal sections of the brain stem, the right column being immunostained for ChAT-positive neurons within the laterodorsal tegmental nucleus (LDT) following unilateral direct injection of UII-saporin (UII-SAP) into the right mesopontine tegmentum (MPT). Scale bar = 200 μm **(B)** Direct injection bilateral of UII-SAP into the MPT reduces the number of ChAT-positive neurons within the LDT compared with Blank-SAP injections (*P* < 0.0001). **(C)** Intraventricular injection of UII-SAP reduced the number of ChAT-positive neurons within the LDT compared to control unconjugated saporin injections (Blank-SAP) (*P* = 0.0003). **(D)** Direct injection of UII-SAP into the LDT does not affect the number of calbindin-positive GABAergic neurons in the LDT compared to Blank-SAP injection (*P* = 0.8857). **(E)** Number of ChAT-positive hypoglossal motor neurons per section following injection of UII-SAP or Blank-SAP is not different (*P* = 0.9470). **(F)** The distance travelled by UII-SAP- and Blank-SAP-injected mice in the centre area of the open field test, which was not different between conditions (*P* = 0.8313). **(G)** Time spent on the Rotarod in three successive trials lasting up to 3 min each is not different between UII-SAP- and Blank-SAP-injected mice. \*\*\**P* < 0.001 Students unpaired two tail t-test; n.s., non-significant. Results are presented as mean ± s.e.m.

To determine the effect of cMPT lesion on the breathing pattern of the mice we used unrestrained whole-body plethysmography during the sleep period. There was a strong trend for increased respiratory rate (**Sup. Fig 2A)**, and the dimensionless measurement of enhanced pause (Penh), scaled relative to the duration of inhalation and exhalation and thus taking breathing variability into account, was reduced by 25% in cMPT-lesioned mice compared to sham-lesioned animals (Blank-SAP, **Fig 2A**). However, no change in tidal volume or minute ventilation was observed during sleep (**Sup. Fig 2B,C**). Nonetheless, when the blood oxygen saturation of the cMPT-lesioned mice at rest was measured for 30 minutes during the sleep period it was found to fluctuate, averaging 80-90%, a significant reduction compared with the >95% oxygen saturation recorded in control mice (**Fig 2 B,C**). These findings indicate that the cMPT lesion affects breathing patterns, causing mild hypoxemia during sleep.

**Figure 2.**
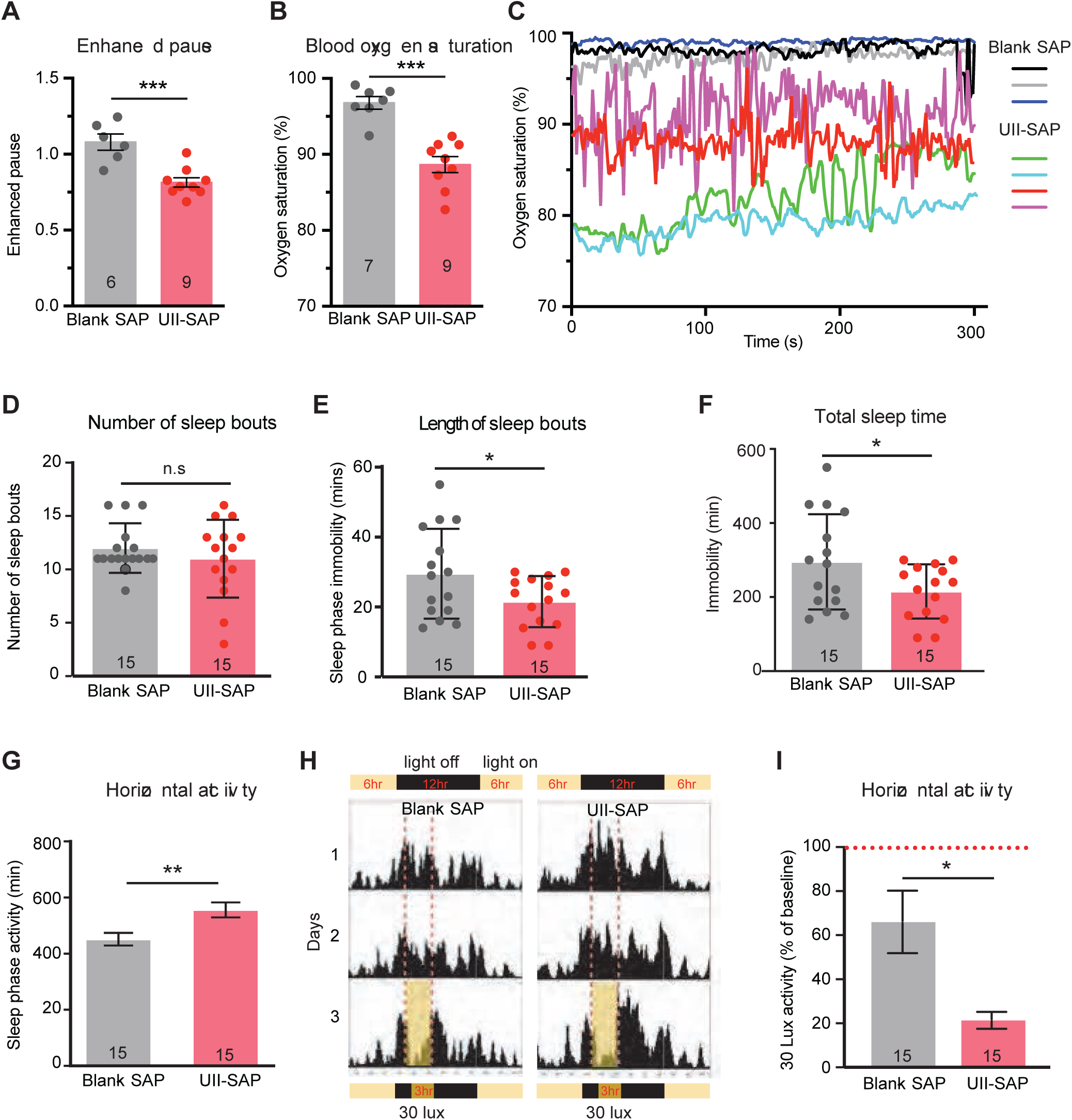
Urotensin II-saporin treatment affects the breathing pattern and induces hypoxia. **(A)** Measure of enhanced pause of mice injected with the UII-SAP or control Blank-SAP recorded by whole body plethysmography during the sleep period (*P* = 0.0005, Student’s unpaired t-test) **(B)** Average blood oxygen saturation level of unrestrained UII-SAP- and Blank-SAP-injected mice measured during their sleep period (*P* < 0.0001, Student’s unpaired t-test). **(C)** Representative traces of blood oxygen saturation levels of mice injected with either UII-SAP or control Blank-SAP. **(D)** Average number of sleep bouts (periods spent inactive lasting at least 10 minutes) during the 12 hour light phase for the first 48 hours of activity recording. (*P* = 0.5023, Paired t-test). **(E)** Average length of sleep bouts (time spent inactive) during the 12 hour light phase in the first 48 hours. (*P* = 0.0475, paired t-test). **(F)** Total time spent inactive during the 12 hour light (sleep) phase in the first 48 hours. (*P* = 0.0475, paired t-test). **(G)** Total time spent inactive during the 12 hour light phase in the first 48 hours (*P* = 0.0014, paired t-test). **(H)** Activity traces of a control and lesioned mouse over 3 days of in 12:12hr light:dark cycle. On day 3 during the dark phase, mice were exposed to 3 hours of 30 lux dim light. **(I)** Time spent active during the 3 hr low light period during the 12 hour light phase on day 3 (*P* = 0.0286, two-tailed Mann Whitney test). \**P* < 0.05; \*\**P* < 0.01; \*\*\**P* < 0.001; Results are presented as mean ± s.e.m.

To determine whether the loss of cMPT neurons altered the sleep patterns of the lesioned mice, animals were individually housed to 12:12 hour light:dark cycle in a PhenoMaster cabinet and the activity was quantified using infrared motion sensors. Lesioned mice had an equivalent number of sleep bouts to control mice (**Fig 2D**) but had shorter sleep bout, significantly less total sleep and were more active during the sleep phase (**Fig 2E,F,G**), indicative of disrupted sleep. To further assess the sleep quality in the mice, a 3 hour dim light pulse (30 lux) was applied during the dark (awake) phase on day 3 to induce sleep (Zhao et al., 2017). The lesioned mice also displayed a sleep debt, showing a greater tendency to sleep during a low light period during their active phase on day 3 (**Fig 2H,I**). Together these phenotypes are consistent with the cMPT lesion inducing OSA in mice.

### cMPT lesion exacerbates the cognitive and pathological features of AD

To determine whether the cMPT lesion and associated OSA phenotypes exacerbated the core features of AD (cognitive impairment, amyloid plaques, neuroinflammation, and neurodegeneration), we used the commonly studied APP/PS1 familial AD mouse model. This strain expresses mutant amyloid precursor protein (APPswe) and presenilin 1 (PS1dE9), leading to overproduction of Aβ, which aggregates and accumulates into amyloid plaques from around 6 months of age, resulting in cognitive impairment (Jankowsky et al., 2004).

We lesioned the cMPT neurons of 8 month old APP/PS1 and tested their cognitive function two months later. No difference was observed between 10 month old lesioned and control groups (sham-lesioned APP/PS1 mice) in terms of time spent in the center of an open field (cMPT lesioned = 46.45 ± 3.66 sec, sham lesioned = 47.88± 5.45 sec). However, cMPT-lesioned APP/PS1 mice displayed severe impairment in hippocampal-dependent learning and memory compared to age-matched unlesioned APP/PS1 mice. In the Y maze, sham-lesioned mice spent significantly more time in the novel arm of the apparatus than in the familiar arm, whereas cMPT-lesioned mice showed no preference (**Fig 3A**). In the Morris water maze, APP/PS1 sham-lesioned mice learned the location of the platform during training, whereas the cMPT-lesioned mice did not (day 1 vs day 5: Blank-SAP, *P* = 0.0039; UII-SAP, *P* > 0.9999; **Fig 3B**). In the probe trial both lesioned and sham-lesioned APP/PS1 mice had similar poor performance (**Fig 3C**), likely reflecting memory impairment due to the animals’ age and genotype, irrespective of lesion status. Similarly, in the active place avoidance task, neither lesioned nor control APP/PS1 mice showed significant learning during 4 days of training (Blank-SAP: day 1 vs day 4, *P* = 0.2286; UII-SAP: day 1 vs day 4, *P* = 0.9999; **Fig 3D**). However, lesioned APP/PS1 mice exhibited a greater impairment in both the learning (**Fig 3D**) and memory (**Fig 3E**) components of this task compared to controls. Together these data indicate that cMPT neuron lesions exacerbate cognitive deficits in aged APP/PS1 mice.

**Figure 3.**
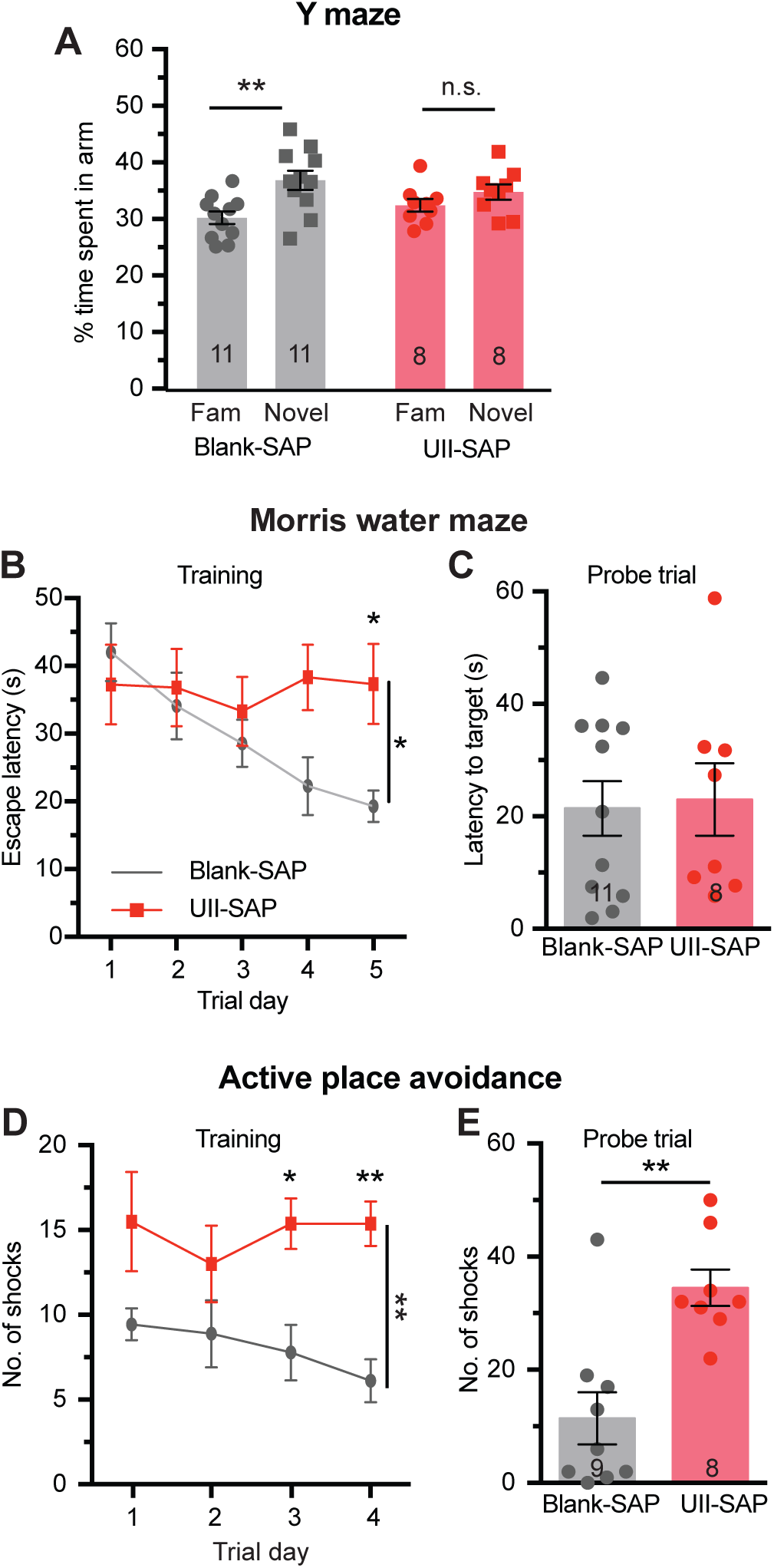
cMPT lesion exacerbates cognitive impairment in the APP/PS1 animal model. **(A)** The percentage time spent in the novel arm of the Y maze on test, compared to the familiar (Fam) arm. Blank-SAP-injected mice displayed a preference for the novel arm, whereas cMPT-lesioned mice had no preference (*P* = 0.0040, two-way ANOVA, Tukey’s multiple comparison test; Blank-SAP: *P* = 0.0072, UII-SAP: *P* = 0.6994). **(B)** Escape latency in the training phase of the Morris water maze. cMPT-lesioned APP/PS1 mice spent significantly more time finding the escape platform on the last two training days (*P* = 0.0150, two-way ANOVA, Tukey’s multiple comparison test; day 5: *P* = 0.0376). **(C)** Latency to reach the platform position in the probe test of the Morris water maze. There were no significant differences in escape latency between groups (*P* = 0.8430, Student’s unpaired t-test). **(D)** The number of shocks received by mice each training day of the active place avoidance test. cMPT-lesioned APP/PS1 mice received significantly more shocks on the last two training days (*P* = 0.0044, two-way ANOVA, Tukey’s multiple comparison test; day3, *P* = 0.0159; day 4, *P* = 0.0022). **(E)** The number of “shocks” the mice received in the probe trial of the active place avoidance test was significantly higher for cMPT-lesioned APP/PS1 mice than non-lesioned APP/PS1 mice (*P* = 0.0012, Student’s unpaired t-test). \**P* < 0.05; \*\**P* < 0.01; n.s., non-significant. Results are presented as mean ± s.e.m.

To determine whether the cMPT-lesioned APP/PS1 mice exacerbate the pathological features of AD, we assessed the number of Aβ plaques, as well as the levels of inflammation and neurodegeneration of the cBF neurons (**Fig 4A**). As expected, the APP/PS1 mice injected with UII-SAP at 8 months of age and sacrificed 3 months later displayed significant cLDT neuronal loss (**Fig 4B**), equivalent to that of cMPT-lesioned wildtype mice (**Fig 1B**). cMPT-lesioned APP/PS1 mice had significantly increased levels of soluble Aβ_42_ in hippocampal lysates (**Fig 4C**), as well as more thioflavin S-positive (ThS) and 6E10-Aβ plaques in the cortex (**Fig 4A,D,E**), compared to sham-lesioned APP/PS1 mice. The levels of microglia (**Fig 4A,F,G**) and activated astrocytes (**Fig 4A,H,I**) in the cortex of MPT-lesioned APP/PS1 mice were also significantly higher. Furthermore, the number of cBF neurons, a feature of AD that occurs early in the human disease but is not readily observed in the familial mouse models (Hampel et al., 2019), was significantly reduced in cMPT-lesioned APP/PS1 mice compared to sham-lesioned APP/PS1 mice (**Fig 4J**).

**Figure 4.**
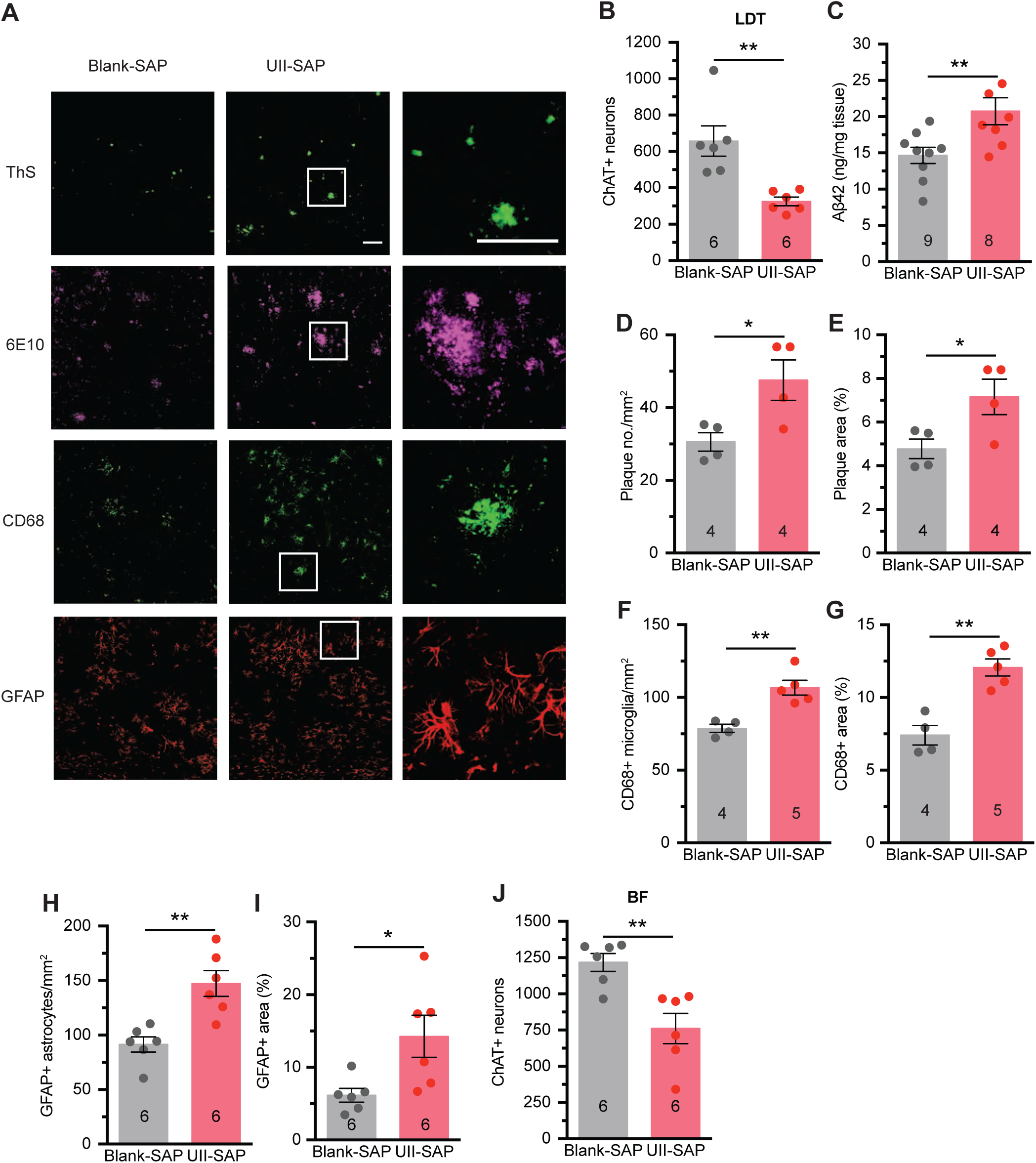
cMPT lesion exacerbates major AD hallmarks of lesioned APP/PS1 mice. **(A)** Representative photomicrographs of sagittal brain sections of hippocampus and cortex of APP/PS1 mice treated with UII-saporin or control Blank-saporin and stained with thioflavin S (ThS), 6E10 (both for Aβ plaques), CD68 and GFAP. Scale bars= 100 μm. **(B)** Average number of ChAT-positive neurons in the LDT of UII-SAP-injected APP/PS1 mice was lower than that of APP/PS1 mice injected with Blank-SAP (*P* = 0.0033). **(C)** Amount of soluble Aβ in the hippocampal lysates of APP/PS1 mice treated with UII-SAP as measured by ELISA was higher than that of APP/PS1 mice injected with Blank-SAP (*P* = 0.0116). Density **(D**; *P* = 0.0320) and area (**E**; *P* = 0.0432) of thioflavin-S-positive Aβ plaque in neocortex of APP/PS1 mice treated with UII-SAP was higher than that of APP/PS1 mice injected with Blank-SAP. Density (**F**; *P* = 0.0028) and area (**G**; *P* = 0.0012) of CD68 immmunopositive staining in the neocortex of the APP/PS1 mice treated with UII-SAP was higher than that of APP/PS1 mice injected with Blank-SAP. Density (**H**; *P* = 0.0023) and area (**I**; *P* = 0.0241) of GFAP immunostaining in the neocortex of the mice treated with UII-SAP was higher than that of APP/PS1 mice injected with Blank-SAP. **(J)** Average number of ChAT-positive basal forebrain (BF) neurons in APP/PS1 mice treated with UII-SAP was lower than that of APP/PS1 mice injected with Blank-SAP (*P* = 0.0038). \**P* < 0.05; \*\**P* < 0.01. Student’s unpaired t-test for panels B-J. Results are presented as mean ± s.e.m.

Together these results indicate that cMPT lesioning causing hypoxemia and sleep disruption, when coupled with a genetic drive to overproduce Aβ can exacerbate the key pathological and cognitive features of AD in mice.

### cBF degeneration is induced by cMPT lesion

Cholinergic basal forebrain (cBF) neurons project to the entire neocortex, regulating cognitive processes including spatial navigation and associative learning (Ballinger et al., 2016). These neurons are among the first to degenerate in AD, with their loss being evident in mild cognitive impairment and idiopathic AD, as well as in individuals who are subsequently diagnosed with cognitive impairment (Hampel et al., 2019). In humans, Aβ load correlates with basal forebrain atrophy and dysfunction (Grothe et al., 2014a; Kerbler et al., 2015a; Chiesa et al., 2019), but it is unclear whether cBF degeneration is a cause and/or effect of Aβ accumulation. In mouse models, oligomeric Aβ can induce cBF degeneration (Sotthibundhu et al., 2008; Knowles et al., 2009), whereas cBF degeneration can exacerbate Aβ accumulation (Gil-Bea et al., 2012; Laursen et al., 2014; Turnbull et al., 2018).

To determine the cause of cBF neuron loss observed in the APP/PS1 mice, we examined the effect of the cMPT lesioning on cBF neuron survival in wildtype (C57Bl6J) mice over a 7 week period. Significant loss of cLDT neurons was evident by 2 weeks, consistent with the nature of the saporin toxin-induced cell death (Hamlin et al., 2013), with more than 50% loss of cLDT cells observed at later time points (**Fig 5A**). The loss of cBF neurons occurred subsequent to the cMPT degeneration between 2 and 4 weeks after surgery (**Fig 5B**), resulting in ∼30% loss of cells from the medial septum, as well as the vertical and horizontal diagonal bands of Broca (**Sup. Fig 3A-C**).

**Figure 5.**
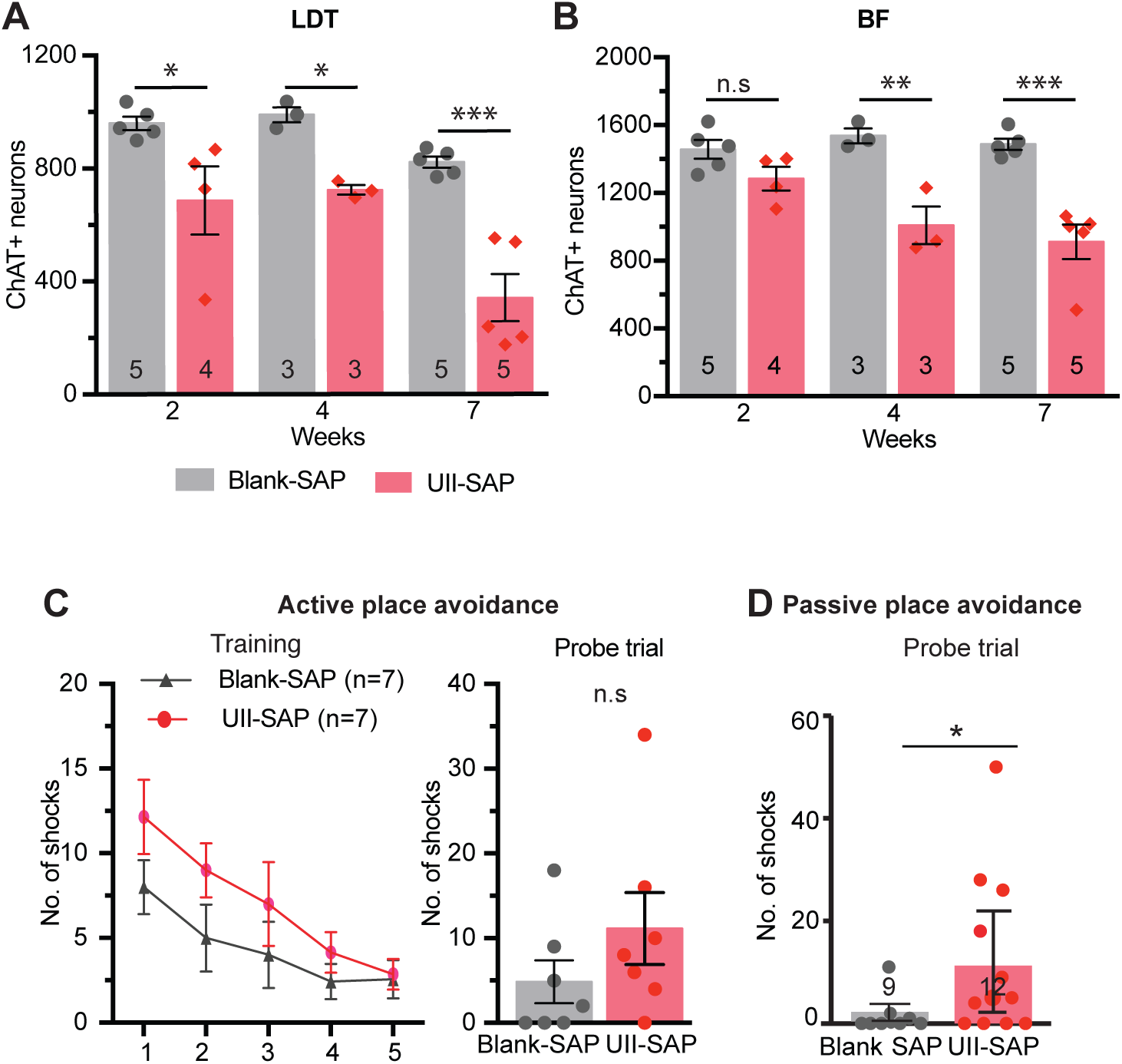
cMPT lesions induce cBF neuronal degeneration and cognitive impairment. (A) The number of ChAT-positive neurons in the LDT of C57Bl6 mice following injection of either UII-SAP or Blank-SAP 2-7 weeks earlier (P < 0.0001, two-way ANOVA, Bonferroni’s multiple comparisons test, 2wk: *P* = 0.0133, 4wk: *P* = 0.0248, 7wk: *P* < 0.0001). (B) The number of ChAT-positive neurons in the basal forebrain (BF) of C57Bl6 mice following injection of either UII-SAP or Blank-SAP 2 to 7 weeks earlier. cBF neuron loss was subsequent to the loss in the LDT (P < 0.0001, two-way ANOVA, Bonferroni’s multiple comparisons test, 2wk: *P* = 0.3142, 4wk: *P* = 0.0010, 7wk: *P* < 0.0001). (C) In the active place avoidance test, the number of shocks was not significantly different between conditions on any given training day (*P* = 0.7580, two-way ANOVA, Tukey’s multiple comparison test) or on the probe test between the cMPT-lesioned mice and sham-lesioned mice (*P* = 0.2279, Student’s unpaired t-test), although a trend for poorer performance was seen. (D) The number of ‘shocks’ received (*P* = 0.04, Student’s unpaired t-test) in the probe trial of the passive place avoidance test was significantly different between the UII-SAP and control blank-SAP groups. \**P* < 0.05; \*\**P* < 0.01; n.s., non-significant. Results are presented as mean ± s.e.m.

Furthermore, the reduction in the number of cBF neurons significantly correlated with the size of the cMPT lesion (r = 0.5746, *P* = 0.0080, R_2_ = 0.3302). No change was observed in the number of parvalbumin-positive GABAergic neurons in the basal forebrain (**Sup. Fig 3C**). This cBF degeneration was also sufficient to cause impairment in visuo-spatial memory consolidation. No significant deficits in the Y maze (**Sup. Fig 3D**) or novel object recognition tasks (**Sup. Fig 3E**), were displayed by cMPT-lesioned wildtype mice or unlesioned mice. However, in the active place avoidance navigation task many of the cMPT-lesioned mice received more shocks during the training and probe trial than control animals (**Fig 5C**). Moreover, in the passive place avoidance task, which requires cBF function (Hamlin et al., 2013), cMPT-lesioned mice displayed a significant memory deficit (**Fig 5D**).

### Sleep deprivation does not result in cBF neuronal loss or Aβ accumulation

To determine whether the disrupted sleep or hypoxia induced by OSA was responsible for causing cBF degeneration and Aβ accumulation, wildtype and aged APP/PS1 mice were subjected to sleep deprivation using a modified ‘flowerpot’ method (**Sup. Fig 4A**), whereby mice entering REM sleep (accompanied by muscle relaxation) are woken by contact with water, preventing sustained sleep. Cohorts of unlesioned mice were placed in sleep-deprivation cages for 20 hours a day, returning to their home cage for 4 hours a day during their sleep phase. The mice became visually sleepy over the course of the experiment, but maintained their body weight (**Sup. Fig 4B,C**). After 4 weeks of sleep deprivation, APP/PS1 animals displayed deficits in Y maze performance that were not exhibited by age-matched APP/PS1 mice housed in equivalent conditions but in cages that lacked water (**Fig 6A**). However, contrary to expectation, latency to first sleep at the end of the paradigm was driven by age rather than cage type (two-way ANOVA, age: *P* = 0.0113 cage: *P* = 0.2392), with sleep deprivation causing long-lasting effects on the animals’ sleep patterns. Sleep-deprived mice had normal diurnal sleep/wake rhythms but a longer sleep latency and a reduction in total sleep time in the 3 days after the paradigm ended (**Sup. Fig 4D-F**). Nonetheless, the number of cBF neurons in the sleep-deprived mice was not significantly different from the number in control mice (**Fig 6B**). The levels of soluble and plaque Aβ (**Fig 6C,D**) and degree of inflammation (**Sup. Fig 5A-D**) in the APP/PS1 mice were also unaltered by the sleep deprivation conditions. This indicates that OSA-induced hypoxia rather than sleep deprivation likely causes the cBF degeneration and exacerbates Aβ accumulation and inflammation.

**Figure 6.**
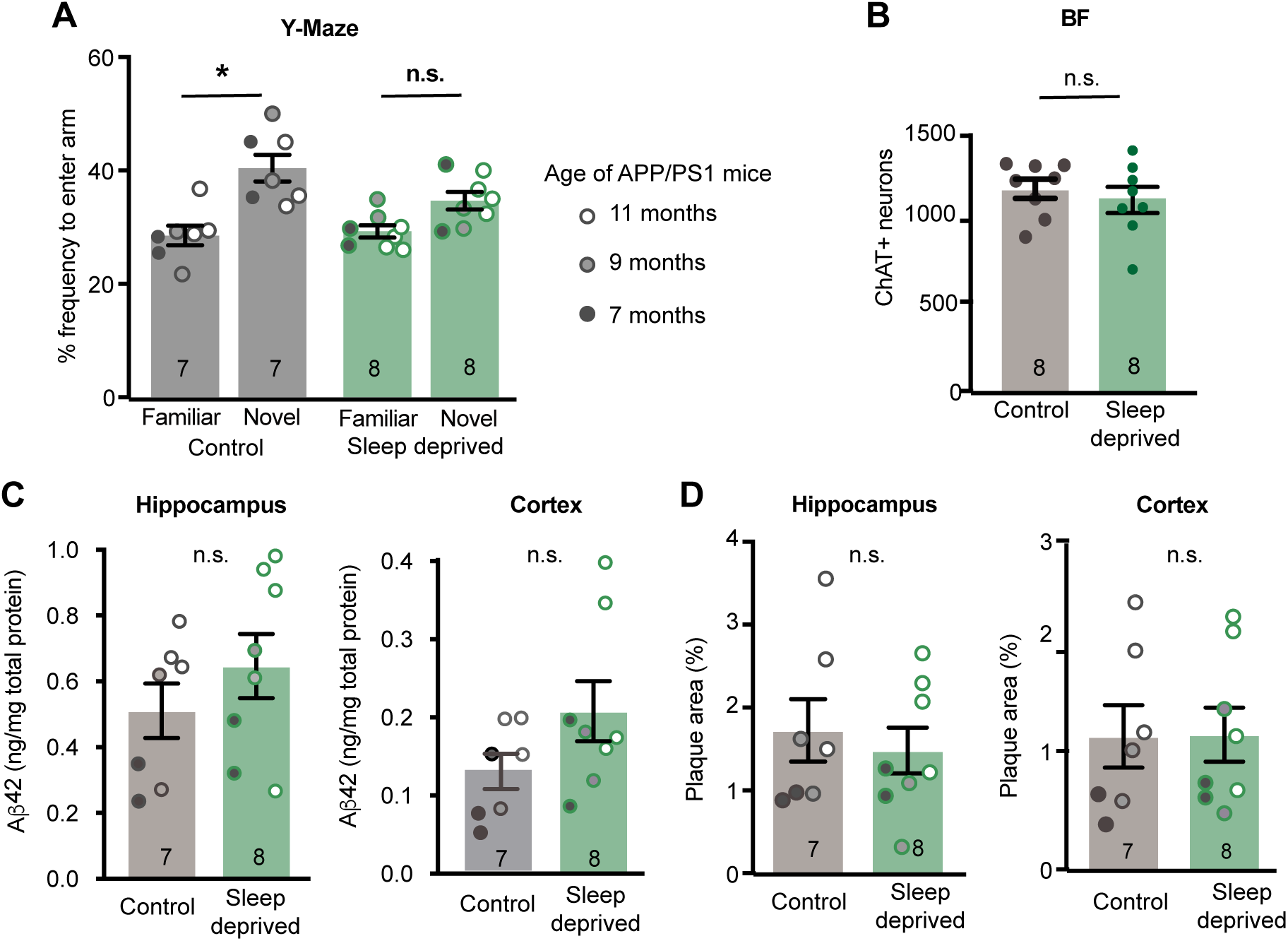
Sleep deprivation causes cognitive impairment but not AD pathology. **(A)** The percentage of time spent in the novel arm of the Y maze on test, compared to the familiar arm. APP/PS1 mice displayed a preference for the novel arm, whereas sleep-deprived mice had no preference (*P* = 0.0477, control: *P* = 0.0050, Sleep-deprived: *P* = 0.1733). **(B)** The number of cBF neurons in C57Bl6 mice following sleep deprivation was equivalent to that of control mice (*P* = 0.624, Student’s unpaired t-test) **(C)** The amount of soluble Aβ in hippocampal (age: *P*= 0.0602, cage: *P =* 0.774, interaction: *P* = 0.8960) and cortical (age: *P* = 0.0733, cage: *P* = 0.155, interaction *P* = 0.734) lysates from as measured by ELISA was not altered by sleep deprivation. **(D)** The area of thioflavin-S-positive Aβ plaque in the hippocampus (age: *P* = 0.0330, cage: *P =* 0.9852, interaction: *P* = 0.6177) and cortex (age: *P* = 0.0128, cage: *P* = 0.410, interaction *P* = 0.666) of APP/PS1 mice was affected by age but not by sleep deprivation. \**P* < 0.05; n.s., non-significant. two-way ANOVA, Sidak’s multiple comparisons for panels A, C and D. Results are presented as mean of pooled age groups ± s.e.m.

### HIF1α and p75_NTR_ mediate cBF death following cMPT lesioning

To test whether the cBF neurons were dying due to exposure to chronic hypoxic conditions, wildtype mice were exposed to reduced (80%) oxygen conditions for 8 hours a day during their sleep period for 4 weeks. At the end of this period, the number of cBF neurons in mice subjected to daily chronic hypoxia was not significantly different from that of mice housed continuously in normoxia (**Fig 7A**), indicating that chronic hypoxia was unable to trigger the same degenerative pathways that were induced by cMPT lesioning. In contrast, chronic intermittent hypoxia (with oxygenation of the air fluctuating from 0 to 100% every 90 seconds) has been reported to induce cBF neuronal degeneration (Row et al., 2007).

**Figure 7.**
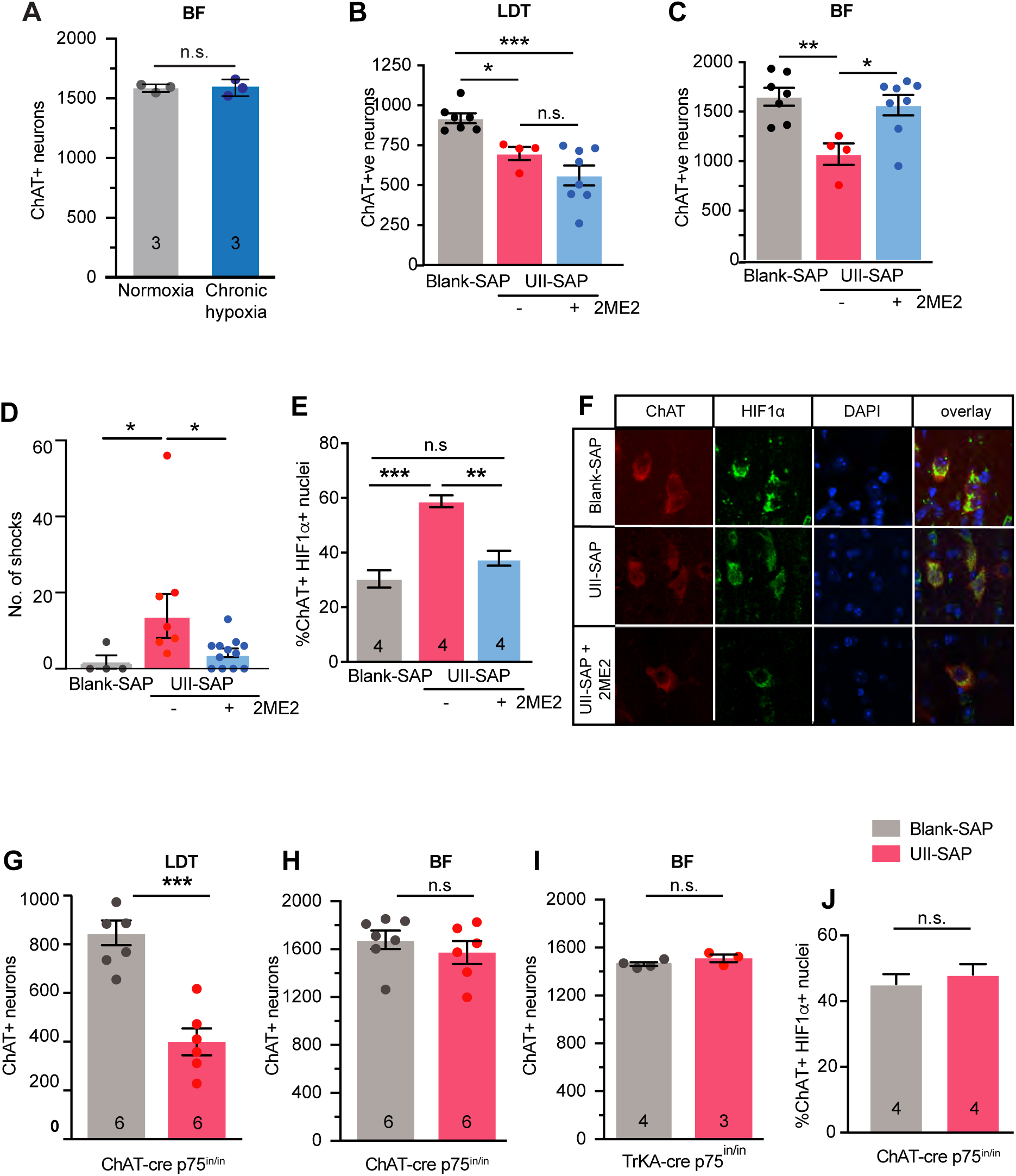
cBF neuronal loss following cMPT lesion is induced by intermittent hypoxia. **(A)** The number of cBF neurons in C67Bl6 mice following 4 weeks of daily sleep-time exposure to hypoxic conditions. The number of cLDT **(B)** and cBF **(C)** neurons in mice following injection of either Blank-SAP (gray bar) or UII-SAP and treated with daily 15mg/kg 2ME2 (blue bar) or vehicle (red bar) for 3 weeks (cBF results: Blank-SAP vs. UII-SAP: *P* = 0.0001, Blank-SAP vs. 2ME2 treated: *P* = 0.0020, UII-SAP vs. 2ME2 treated: *P* = 0.5401). 2ME2 treatment protects cBF neurons from the effects of lesioning. **(D)** Performance of 2ME2-treated (blue bar) and untreated UII-SAP-lesioned (red bar) mice compared to Blank-SAP-lesioned mice (gray bar) in the test phase of the passive place avoidance task. (Blank-SAP vs. UII-SAP: *P* = 0.0152, Blank-SAP vs. 2ME2 treated: *P* = 0.4014, UII-SAP vs. 2ME2 treated: *P* = 0.0112) **(E)** The percentage of cBF neurons in which HIF1α immunostaining was present in the nucleus in Blank-SAP (gray bar) or UII-SAP mice treated with daily 15mg/kg 2ME2 (blue bar) or vehicle (red bar) for 3 weeks. (Blank-SAP vs. UII-SAP: *P* = 0.0001, Blank-SAP vs. 2ME2 treated: *P* = 0.1766, UII-SAP vs. 2ME2 treated: *P* = 0.0011) **(F)** Representative confocal images of basal forebrain sections immunostained for ChAT (red) and HIF1α (green) and nuclei (DAPI; blue). **(G)** The number of LDT neurons in UII-SAP-injected ChAT-cre p75_in/in_ mice was decreased compared with that of Blank-SAP-injected ChAT-cre p75_in/in_ mice (*P* <0.0001). **(H)** The number of cBF neurons in UII-SAP-injected ChAT-cre p75_in/in_ mice was not significantly different from that in Blank-SAP-injected ChAT-cre p75_in/in_ mice (*P* = 0.408). **(I)** The number of cBF neurons in UII-SAP-injected TrkA-cre p75_in/in_ mice was not significantly different from that in Blank-SAP-injected TrkA-cre p75_in/in_ mice (*P* = 0.2006). **(J)** The percentage of cBF neurons in which HIF1α immunostaining was present in the nucleus in Blank-SAP or UII-SAP injected ChAT-cre p75_in/in_ mice after 4 weeks. (Blank-SAP vs. UII-SAP: *P* = 0.0001, Blank-SAP vs. 2ME2 treated: *P* = 0.1766, UII-SAP vs. 2ME2 treated: *P* = 0.0011) \**P* < 0.05; \*\**P* < 0.01; \*\*\**P* < 0.001; n.s., non-significant. one-way ANOVA Tukey’s multiple comparison test for panels B-E and Students unpaired t-test for panels A,G-J. Results are presented as mean ± s.e.m.

To test this theory, cMPT-lesioned mice were treated for 4 weeks with the drug 2-methoxyestradiol (2ME2). This compound binds to and prevents the nuclear translocation of hypoxia-inducible factor 1 alpha (HIF1α), the expression of which is induced by low oxygen conditions and mediates the transcription of genes such as vascular endothelin growth factor (VEGF) to improve cerebral blood flow and oppose the toxicity of hypoxia. However, prolonged or persistent activation of HIF1α has also been linked to cell death mediated by pro-apoptotic Bcl-2 family members e.g. via BNIP3 (Dayan et al., 2006; Iyalomhe et al., 2017). Two weeks after cMPT lesioning, wildtype mice were injected daily for 3 or 4 weeks with 2ME2. The extent of cMPT neuron loss was equivalent between 2ME2-treated and untreated lesioned mice, and significantly reduced compared to the cMPT number in sham-lesioned control animals (**Fig 7B; Sup. Fig 6B**). However, the number of cBF neurons in the 2ME2-treated lesioned animals was significantly increased compared to that of untreated lesioned animals, and similar to that of sham-lesioned mice (**Fig 7C; Sup. Fig 6C**). The 2ME2-treated lesioned animals also exhibited a degree of improved memory performance in the cBF-dependent passive place avoidance task (**Fig 7D**). Furthermore, the percentage of cBF neurons with HIF1α present in the nucleus (indicative of its activation) in lesioned, untreated animals was significantly higher than that in both treated cMPT-lesioned mice, and sham-lesioned mice (**Fig 7E,F**). This indicates that 2ME2 engaged its target, and that HIF1α activity mediates cBF death following cMPT lesioning.

cBF neurons express the neural cell death receptor p75 neurotrophin receptor (p75_NTR_) at high levels throughout life. p75_NTR_ regulates cBF neuronal death during development and in disease conditions (Ibanez and Simi, 2012; Boskovic et al., 2014). Furthermore, p75_NTR_ has been reported to regulate the activity of HIF1α, with low oxygen levels causing cleavage of p75_NTR_, which in turn reduces HIF1α degradation *in vitro* (Le Moan et al., 2011; Tong et al., 2018). Therefore, we asked whether cBF neurons lacking p75_NTR_ expression were susceptible to cMPT lesion-induced loss. In ChAT-cre p75_in/in_ mice (Boskovic et al., 2014), in which p75_NTR_ is removed from cholinergic neurons, a cMPT lesion equivalent to that observed in control p75_fl/fl_ mice did not result in any loss of cBF neurons (**Fig 7G,H**). Although p75_NTR_ is not expressed by adult cMPT neurons, its expression would be genetically removed from cMPT neurons in the ChAT-cre p75_in/in_ mice. Therefore, we repeated the experiment in mice in which p75_NTR_ was removed from cBF neurons but not cMPT neurons (TrkA-cre p75_in/in_ mice), again finding no loss of cBF neurons (**Fig 7I**). Furthermore, in both lesioned and sham-lesioned p75_fl/fl_ mice, HIF1α immunostaining did not colocalize in the nucleus in the majority of cBF neurons neurons (**Fig 7J**). These data indicate that the loss of cholinergic cBF neurons following cMPT lesioning is mediated by p75_NTR-_dependent HIF1α nuclear activity.

### Preventing hypoxia prevents AD features after cMPT lesion

The standard treatment for OSA in human sufferers is the use of continuous positive airway pressure (CPAP) therapy during sleep, which physically maintains airway opening and thus prevents blood oxygen desaturation and arousal from sleep. We therefore asked whether treatment of OSA mice during their sleep phase with a high-oxygen environment could protect cBF neurons from cMPT lesion-induced death and/or A^□^ accumulation. We first empirically determined that a high (40%) oxygen environment restored the blood oxygen levels of cMPT-lesioned mice to >95%. Starting 2 weeks after lesion surgery, wildtype (**Fig 8A-C**) and aged APP/PS1 (**Fig 8D-K**) mice were then treated daily with high oxygen during their 12 hour sleep period for either 2 weeks (*i.e.* weeks 2-4 after cMPT lesioning) or 4 weeks (weeks 2-6), respectively. As expected, high oxygen treatment had no effect on the extent of the cMPT lesion in both wildtype (**Fig 8A**) and APP/PS1 mice (**Fig 8D**). Nonetheless, the treatment restored sleep patterns, and prevented the development of a sleep debt (**Fig 8C**), demonstrating that these phenotypes were not induced directly by the cMTP lesioning, further validating our OSA model. Furthermore, the number of cBF neurons in both wildtype (**Fig 8B**) and APP/PS1 (**Fig 8E**) high oxygen-treated cMPT-lesioned animals was significantly higher than that of untreated lesioned mice, being similar to that of sham-lesioned animals in standard housing (**Fig 8B**), confirming that the cBF degeneration was a result of the OSA phenotype. Moreover, in the lesioned APP/PS1 cohort, the degree of amyloid pathology **(Fig 8F,G; Sup Fig 7**) and inflammation (**Fig 8H-K; Sup Fig 7**) were significantly reduced by the daily high-oxygen treatment.

**Figure 8.**
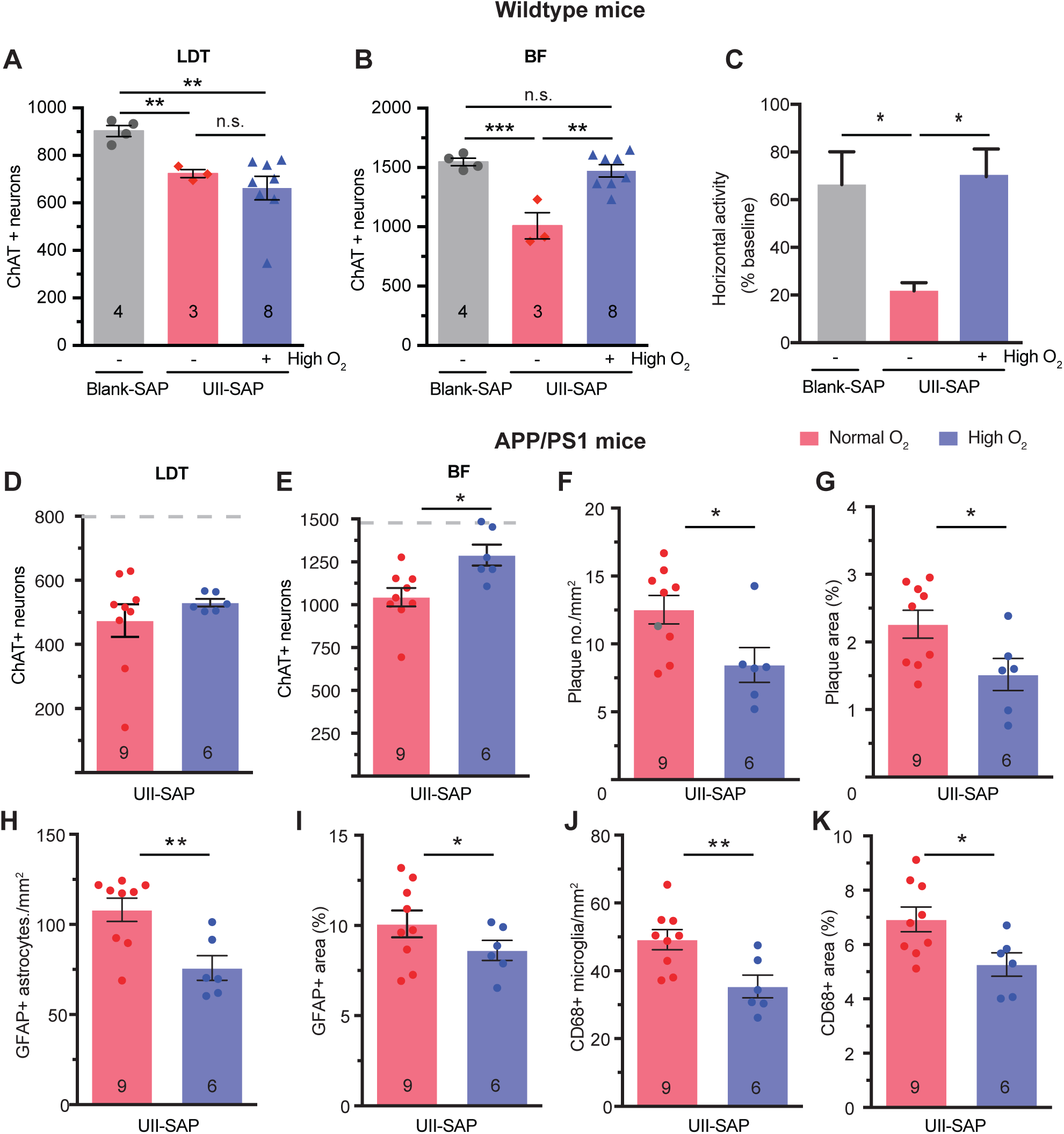
High oxygen treatment protects from OSA-exacerbated AD phenotypes. **(A)** Number of cMPT neurons in wildtype Blank-SAP- (grey bar) and UII-SAP-injected mice. Both high O2-treated (purple bar) and normoxia UII-SAP injected (red bar) mice had a significant loss of cMPT neurons compared to Blank-SAP-injected mice (*P* = 0.0137, one-way ANOVA, Tukey’s multiple comparison test; Blank-SAP vs UII-SAP -O_2_: *P* = 0.0021, Blank-SAP vs UII-SAP +O_2_: *P* = 0.0089, UII-SAP -O_2_ vs +O_2_: *P* = 0.4929). **(B)** The cBF neuronal number of UII-SAP-injected mice subjected to high oxygen (purple bar) was significantly higher than that of mice subjected to normoxia (red bar), and not significantly different from that of Blank-SAP controls (gray bar), whereas the number of cBF neurons in UII-SAP-injected normoxia-treated mice (red bar) was significantly reduced compared with that of Blank-SAP controls (gray bar) (*P* = 0.0006, Blank-SAP vs UII-SAP -O_2_: *P* = 0.0009, Blank-SAP vs UII-SAP +O_2_: *P* = 0.6790, UII-SAP -O_2_ vs +O_2_: *P* = 0.0011). **(C)** Mice placed in 40% oxygenated (high O_2_) for 8 hours a day during the sleep period for 2 weeks starting 2 weeks after injection with UII-SAP (purple bar) do not have a sleep debt compared to UII-SAP-injected mice subjected to normoxia (data from Fig 2H). (*P* = 0.04) Number of cLDT **(E)** and cBF **(F)** neurons in APP/PS1 mice injected with UII-SAP and either untreated (normoxia, normal oxygen) or treated daily with high oxygen. Number **(G**; *P* = 0.0320) and area (**H**; *P* = 0.0432) of thioflavin-S-positive Aβ plaques in the neocortex of APP/PS1 mice injected with UII-SAP and treated with normoxia or high oxygen. Density **(I**; *P* = 0.0056) and area (**J**; *P* = 0.0432) of GFAP-positive microglia in the neocortex of APP/PS1 mice injected with UII-SAP and treated with normoxia or high oxygen. Density **(K**; *P* = 0.0098) and area (**L**; *P* = 0.0264) of CD68-positive astrocytes in the neocortex of APP/PS1 mice injected with UII-SAP and treated with normoxia or high oxygen. \**P* < 0.05; \*\**P* < 0.01; \*\*\**P* < 0.001; n.s., non-significant. One way ANOVA with Tukey’s multiple comparison test for panels **A**-**C**, Student’s unpaired t-test for panels **D**-**K**. Results are presented as mean ± s.e.m.

## Discussion

Despite strong epidemiological links between OSA and the development of AD, the mechanisms underlying this increased risk are unclear. We developed a naturalistic model of OSA in mice, replicating key features of the human condition without comorbid risk factors such as cardiovascular disease and diabetes. Using our OSA model, we demonstrated that cBF neurodegeneration and cognitive impairment, two early features of AD, are induced by the resultant intermittent hypoxia and, in a genetically susceptible (familial) AD mouse, OSA exacerbates additional pathological features such as plaque load and inflammation. In contrast, although induction of chronic sleep deprivation of mice (for an equivalent period of time to that in which mice were subjected to OSA) impaired working memory, it neither induced cBF neuronal dysfunction nor exacerbated Aβ accumulation. This suggests that intermittent hypoxia is a major mechanism by which OSA, even in the absence of comorbidities, could promote AD in humans.

### Urotensin 2-saporin toxin lesions of cMPT replicates key features of human OSA

Hallmarks of patients with untreated OSA include recurrent nocturnal arousals from sleep, caused by frequent episodes of pharyngeal obstruction. These cause hypopneas (≥4% decrease in arterial oxygen saturation; (Ward et al., 2013)), and hypoxemia (blood oxygen saturation during sleep of <85%) and thus intermittent hypoxia (Rosenzweig et al., 2014), resulting in wakefulness and frequent sleep interruptions. Our model of OSA phenocopies these hallmarks. cMPT-lesioned mice displayed subtle changes to their breathing during sleep, specifically in enhanced pause, a feature of sleep apnoea patients that compensates for the increased upper airway resistance and compliance (Ramirez et al., 2013). Furthermore, the mice exhibited fluctuating hypoxemia, another feature of OSA with hypoxia, comparable to OSA patients with moderate to severe oxygen-desaturation indices. Finally, the cMPT-lesioned mice had disrupted sleep patterns that resulted in a sleep debt, the equivalent of human daytime sleepiness. As prevention of hypoxia during the sleep phase by keeping blood oxygen saturation above 95% prevented the sleep debt, it is reasonable to conclude that the sleep disruption was caused by hypopneas or apnoeas causing waking. Although the breathing changes during sleep are artificially induced, cMPT lesioning therefore represents an authentic animal model of OSA.

### OSA induces pathological hallmarks AD

Untreated elderly patients with OSA exhibit cognitive decline at twice the rate, and develop AD at an earlier age, than the general population (Ancoli-Israel et al., 2008; Osorio et al., 2011; Yaffe et al., 2011). Similarly, in our study, induced OSA in older APP/PS mice (already predisposed to overproduce Aβ peptides) resulted in more severe cognitive impairment in their age-matched littermates, and exacerbated many of the pathological features of AD, including induction of cBF degeneration, Aβ accumulation within the cortex and hippocampus, and increased inflammation represented by astro- and glio-genesis. However, even in the absence of a predisposition to AD pathology, OSA sequelae caused cBF degeneration and cognitive decline. cBF degeneration is associated with mild cognitive impairment in humans (Grothe et al., 2014a) as well as rodents, particularly impairing higher order cognitive functions, including allothetic and egocentric navigation, as measured in place avoidance tasks (Baxter and Chiba, 1999; Hort et al., 2007; Hamlin et al., 2013; Hort et al., 2014; Kerbler et al., 2015b; Turnbull et al., 2018). Although the hippocampal-dependent, spatial memory impairment of cMPT-lesioned APP/PS mice was concurrent with increased Aβ pathology, the coincident cBF neuronal degeneration likely contributed to the behavioural phenotype (Chiesa et al., 2019). These results indicate that OSA even in the absence of other comorbidities can induce or enhance a range of phenotypes associated with AD development. However, as not everyone with OSA develops dementia, it was also important to understand the characteristics of the OSA that increased development of AD phenotypes.

### Intermittent hypoxia underpins the risk of promoting AD features

Sleep deprivation is well known to cause reversible cognitive impairment in humans (Verstraeten and Cluydts, 2004), and the glymphatic system, which can clear Aβ from interstitial fluid and is active during sleep (Tarasoff-Conway et al., 2015; Zuroff et al., 2017), has been proposed as a mechanism by which poor sleep could be a risk for AD. In our study, APP/PS mice subjected to sleep deprivation alone (or to OSA) displayed impairment in Y maze performance. However, sleep deprivation alone was not sufficient to increase Aβ deposition or change Y maze performance in young wildtype animals. We cannot rule out a contribution to AD phenotypes of poor glymphatic clearance or sleep disruption in our OSA model or in patients, particularly if sleep disruption causes persistent changes to sleep levels over a longer period. However, our results suggest that another feature of OSA, intermittent hypoxia, contributes more significantly to the progress of AD pathology than sleep disruption.

We determined that the death of cBF neurons was induced by cMPT lesioning but not by exposure to daily chronic hypoxia. Nonetheless, inhibition of HIF1α activity by 2ME2 or preventing the expression of the low-oxygen-activated and HIF1α regulator, p75_NTR_ (Le Moan et al., 2011; Tong et al., 2018), in cBF neurons prevented their loss in OSA model mice. These results indicate that cBF degeneration induced by OSA is dependent on hypoxia/HIF1α -regulated pathways. Our conclusion is consistent with other reports in which rats exposed to intermittent hypoxia for 14 days (alternating 90s epochs of 21% and 10% O_2_ during sleep) developed deficits in water maze spatial memory performance and reductions in cBF neuron number that mirror the reduction in the OSA model (Row et al., 2007). Similarly, mice exposed to intermittent hypoxia during sleep (alternating 60s epochs of 21% and 0% O_2_ for 8 hours) developed hippocampal changes and cognitive impairment in the Barnes maze spatial memory test (Arias-Cavieres et al., 2019), which is sensitive to cBF function (Greferath et al., 2000; Barrett et al., 2010), and these phenotypes were prevented by heterozygous knockout of HIF1α (Arias-Cavieres et al., 2019).

Why intermittent *versus* chronic hypoxia induce different cBF outcomes may be explained by cell culture experiments in which chronic hypoxia conditions cause HIF1α mRNA levels to be down-regulated after several hours by negative feedback loops, whereas intermittent hypoxia does not active these transcriptional pathways, resulting in increased HIF1α mRNA levels (Cavadas et al., 2017). The extent of HIF1-induced downstream gene expression and repression also varies depending on exposure to chronic or intermittent hypoxia (Cavadas et al., 2017; Martinez et al., 2019). Counterintuitively, chronic hypoxia can also provide cellular protection (Navarrete-Opazo and Mitchell, 2014). We therefore propose that the fluctuations in hypoxemia observed in our OSA model are representative of intermittent hypoxia in the brain, which in turn is responsible for the observed cBF degeneration.

This conclusion is consistent with correlative findings from human populations. The cognitive decline in OSA is linearly correlated with hypoxemia (Adams et al., 2001; Daulatzai, 2014), and a diagnosis of dementia in OSA patients has been associated with the extent of hypoxia rather than sleep disruption (Yaffe et al., 2011; Yaffe et al., 2014). Cognitive impairment is also more common and more severe in OSA patients with hypoxemia (Findley et al., 1986; Incalzi et al., 2004). Additional studies will be required to determine the physiologically relevant tissue-specific levels of hypoxia in our model, and the degree of oxygen volatility required to induce degenerative HIF1a-activated pathways in cBF neurons. Nonetheless, our work highlights intermittent hypoxia as the likely major factor in OSA that causes degenerative changes associated with AD.

### cBF degeneration is an early AD feature that can drive other AD features

Our finding that OSA exacerbates Aβ accumulation in genetically susceptible AD mice is consistent with a previous report that exposure of mice to chronic intermittent hypoxia increases Aβ42 levels in a triple Alzheimer’s disease model mouse (Shiota et al., 2013). Similarly, higher brain Aβ (Spira et al., 2014; Elias et al., 2018), or decreased Aβ and increased phosphorylated tau in the cerebrospinal fluid (Osorio et al., 2014b; Osorio et al., 2014a), has been associated with a higher oxygen desaturation in cognitively impaired OSA subjects. One explanation for these observations could be that hypoxia and HIF1α-mediated transcription can directly up-regulate the expression of β-secretase (Zhang et al., 2007), while HIF1α can operate as a regulatory subunit of γ-secretase during hypoxia (Villa et al., 2014), both of which could increase Aβ production via the amyloidogenic pathway. However, exacerbation of Aβ accumulation in mice that are already overproducing the peptide could equally have occurred through an indirect mechanism, specifically cBF degeneration. Although in this study we did not discern whether the cBF degeneration occurred in parallel with, or exacerbated the other AD pathology, previous research strongly implicates the latter.

cBF neuronal lesioning in a range of mouse AD models worsens many features of AD, including promoting cognitive impairment, and increasing Aβ accumulation, plaque load and inflammation (as observed herein), as well as causing tau hyperphosphorylation, hippocampal degeneration, and reduced BDNF expression (Gil-Bea et al., 2012; Ramos-Rodriguez et al., 2013; Hartig et al., 2014; Laursen et al., 2014; Turnbull et al., 2018). On the other hand, AD model rodents with increased cBF connectivity (Boskovic et al., 2014; Boskovic et al., 2018; Qian et al., 2018) or those which are treated with muscarinic receptor agonists display cognitive improvements and significantly reduced age-dependent A□ accumulation and inflammation compared to control rodents (Fisher et al., 2016). Cholinergic activation of muscarinic receptors also stimulates non-amyloidogenic APP processing (Rossner et al., 1998), thereby regulating the levels of Aβ in the interstitial fluid (Welt et al., 2015). Similarly, in human subjects, reduced basal forebrain volume (as measured by structural MRI, and indicative of cholinergic degeneration) predicts Aβ burden (Grothe et al., 2014a; Teipel et al., 2014; Kerbler et al., 2015a) and is evident in mild cognitive impairment and idiopathic AD cohorts (Grothe et al., 2014a; Grothe et al., 2014b; Daulatzai, 2015), as well as in individuals who are subsequently diagnosed with cognitive impairment (Grothe et al., 2013; Grothe et al., 2014a; Kerbler et al., 2015a; Kerbler et al., 2015b). Furthermore, basal forebrain degeneration, together with the accumulation of Aβ precedes and causes entorhinal cortical degeneration in humans (Schmitz et al., 2016). Therefore, it is probable that the hypoxia-induced death of cBF neurons directly contributed to the other AD phenotypes in our OSA model, and may represent a decisive factor in the aetiology of idiopathic AD. Determining the extent of cBF degeneration in people suffering from intermittent hypoxia as a result of OSA would test this interpretation (Atkins et al., 2016).

Finally, the effect of preventing hypoxemia and sleep disruption in the OSA mice during their daily sleep period, which further validates the OSA model, protected against the development of the phenotypes associated with AD. Similarly, emerging studies indicate that CPAP treatment of people with OSA may reduce neurodegeneration (Owen et al., 2019) and delay the progression of cognitive impairment (Richards et al., 2019), particularly that relating to attention and executive function in which cBF neuronal function is strongly involved (Osorio et al., 2014b; Yaffe et al., 2014). The implication of these studies, together with our findings that the prevention of hypoxemia during the sleep period in cMPT-lesioned mice reduced cBF degeneration and Aβ accumulation, is that early diagnosis and treatment of OSA with CPAP may significantly decrease the longitudinal risk of developing irreversible cognitive impairment. Our study therefore provides a rational impetus to longitudinally study CPAP as a prophylactic treatment to reduce the incidence of AD (Spira and Gottesman, 2017).

### Summary

Our results suggest that OSA, even in the absence of other comorbidities, is a risk factor for the development of AD. Specifically our work highlights intermittent hypoxia as the decisive feature, as it induces cBF neuronal degeneration and cognitive impairment, with a coincident or subsequent increase in the levels of Aβ when linked to a genetic risk for Aβ accumulation (potentially including an ApoE4 genotype; (Yaffe et al., 2014; Wollam et al., 2015)). Although treating the hypoxemia in our OSA model or giving CPAP in humans reduces respiratory disturbance during sleep, and improves working memory, some cognitive aspects such as complex attention and executive function remain impaired (Daulatzai, 2015), daytime sleepiness can continue, and Aβ load can remain unchanged (Elias et al., 2018), all of which might reflect irreversible cBF neuronal loss that predates the commencement of CPAP treatment. Therefore, targeting cholinergic function (through cholinergic esterase inhibition or muscarinic agonists; (Hampel et al., 2019)), cBF health (through p75_NTR_ modifiers or neurotrophic mimetics; (Longo and Massa, 2013)) or specific HIF1α pathways (2ME2 is a current treatment for tumour angiogenesis) represent candidate treatments for preventing AD in the OSA ‘at risk’ population – those with intermittent hypoxia.

## Acknowledgements

This work was supported by the National Health and Medical Research Council of Australia (GNT1049236 to EJC and MCB; GNT 1162505 to EJC), the Brain Foundation, the Mason Foundation and support from the Clem Jones Centre for Ageing Dementia Research. We thank Advance Targeting Systems for their generous supply of UII-SAP and Blank-SAP. Work was performed in the Queensland Brain Institute’s Advanced Microscopy Facility with support from Luke Hammond, the Queensland Brain Institute’s Animal Behavioural Facility with support from Daniel Blackmore, and the School of Biomedical Sciences Integrated Physiology Facility with support from Stuart Mazzone and Melanie Flint. We thank Queensland Brain Institute’s Animal Facility staff particularly Liesel MacDonald, Michael Lardelli (The University of Adelaide, Australia) for helpful suggestions, and Rowan Tweedale for manuscript editing.

## Author contributions

L.Q., O.R., M.C.B. and E.J.C. designed the experiments, L.Q., L.C., N.M., M.M., and A.S. performed experiments, all authors analysed data, discussed the work and wrote or commented on the manuscript.

## Supplementary Material

**Supplementary Figure 1.**
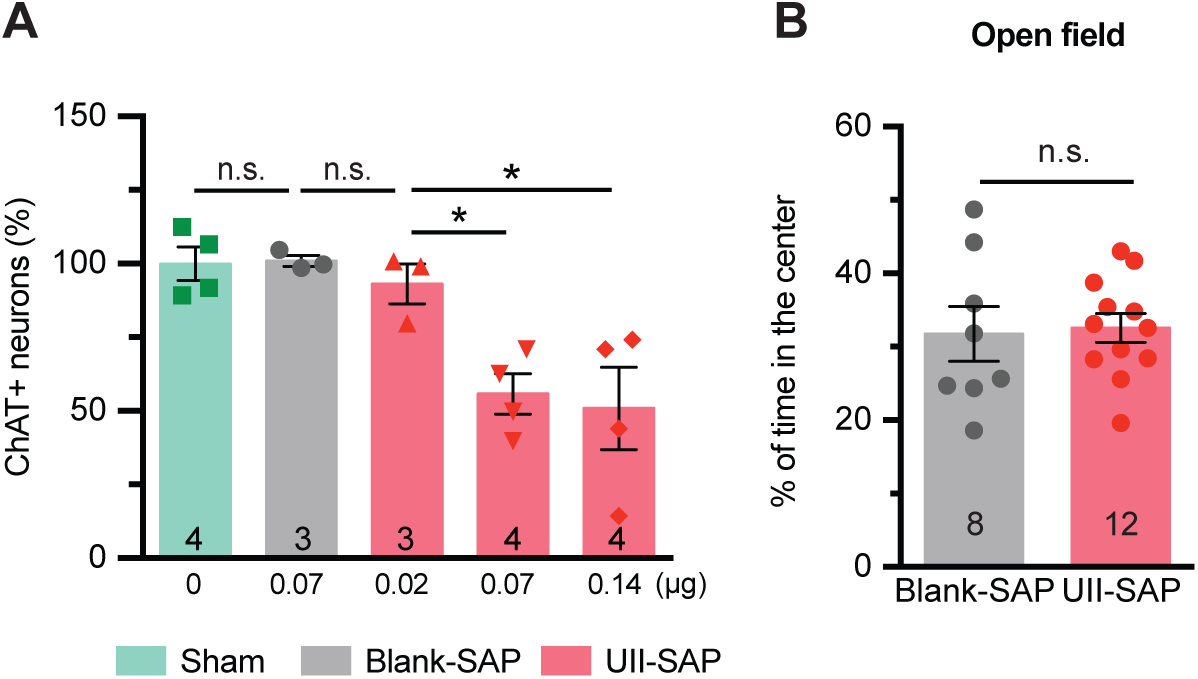
Dose response of UII-SAP-induced cholinergic mesopontine neuronal loss. Four weeks after direct injection with UII-SAP or control, animals were sacrificed and cholinergic neurons in the mesopontine tegmentum were counted. **(A)** Number of ChAT-positive neurons within the LDT following the direct injection of UII-SAP or IgG-SAP into the MPT at the concentrations shown (** *P* = 0.0015, one-way ANOVA, Tukey’s multiple comparison test; Sham vs Blank-SAP: *P* > 0.9999. **(B)** Time spent in the center area in the open field test by cMPT-lesioned and sham-lesioned mice (*P* = 0.5845, Student’s unpaired t-test). Results are presented as mean ± s.e.m.

**Supplementary Figure 2.**
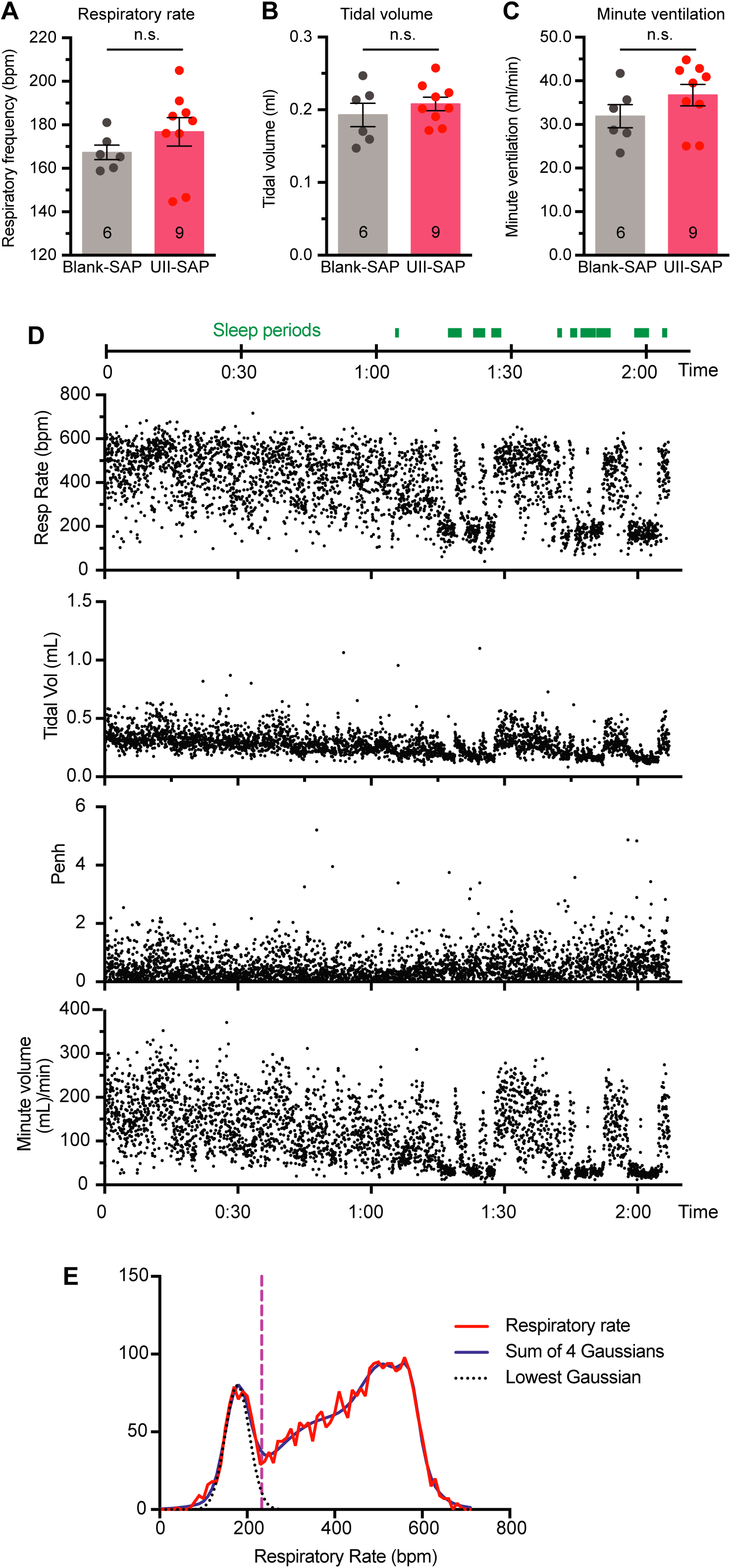
Whole body plethysmographic data analysis. Whole body plethysmography has been used to record respiration during sleep and wake cycles in unrestrained, freely moving mice. In this study measure, mice were excluded from the analysis if they remained active in the chamber and/or their breathing was not characteristically in a sleep stage. Mice were classified as being in a sleep stage if they were inactive with eyes closed, and respiratory rate, tidal volume and minute volume were all reduced and showed low variability for a minimum period of 10 s. Respiratory rate (**A**; *P* = 0.2916), tidal volume (**B;** *P* = 0.3990), and minute ventilation (**C**; *P* =0.2150) of mice injected with the UII-SAP or control Blank-SAP recorded by whole body plethysmography during their quiet sleep period. **(D)** Example of plethysmographic data: sleep periods (green bars) were associated with lower respiratory rate, decreased tidal volume, and decreased minute volume, but not enhanced pause (Penh). **(E)** The binned respiratory rate data (red line), with the fitted sum of 4 Gaussian distributions (blue line) used to define quiet sleep periods. Sleep periods were defined as 5 or more consecutive respiratory rate measurements, where respiratory rate was less than the mean of the lowest Gaussian distribution (dashed black line) + 2 SD of the lowest Gaussian distribution. For this animal, the accepted respiratory rate for quiet sleep was < 233 bpm (vertical purple line).

**Supplementary Figure 3.**
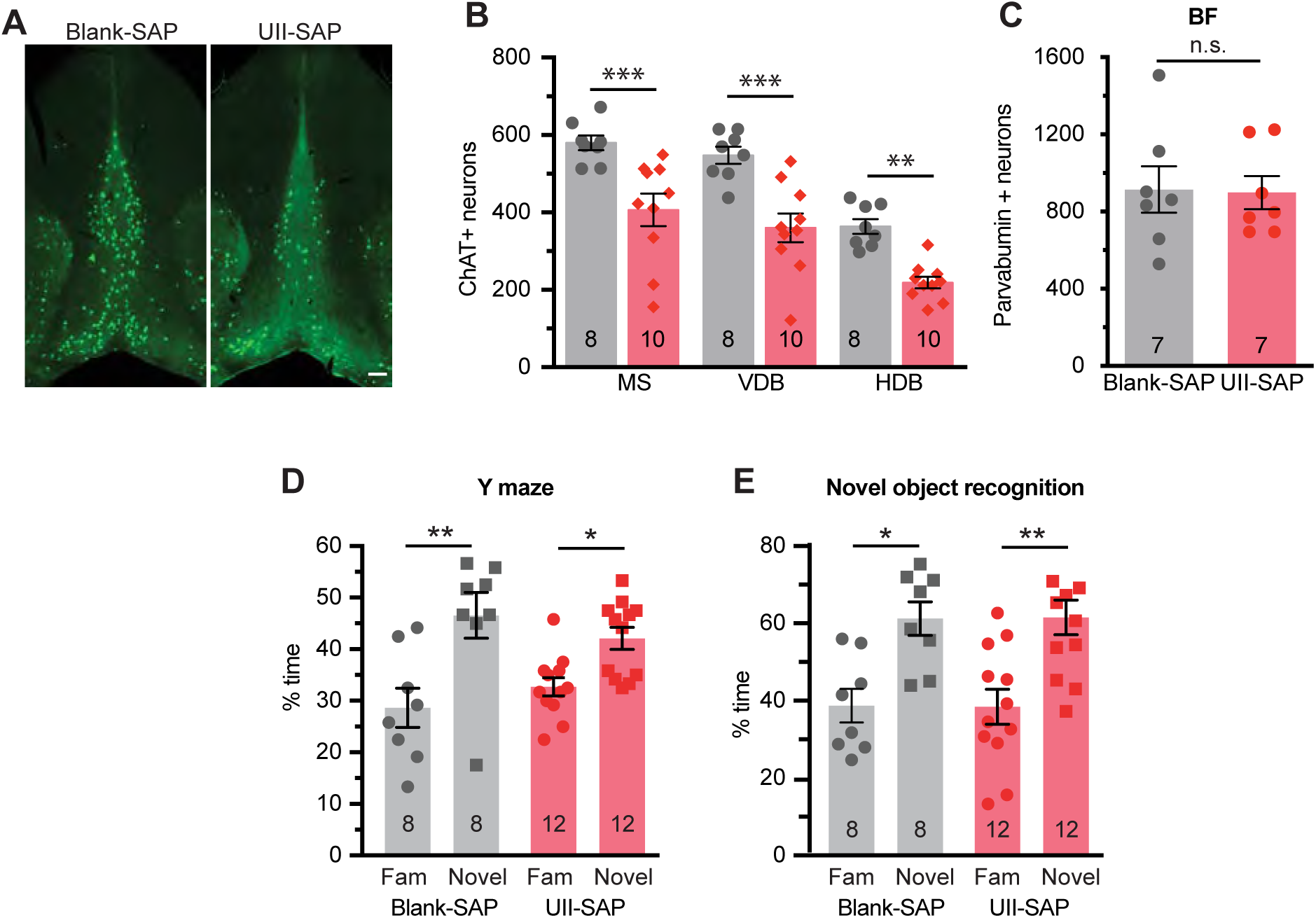
Cholinergic basal forebrain degeneration and behaviour following cMPT lesion. **(A)** Representative photomicrographs of coronal sections of the basal forebrain immunostained for ChAT-positive neurons following the direct injection of UII-SAP or control Blank-SAP) into the MPT. **(B)** The number of ChAT-positive neurons in the medial septum (MS), the vertical diagonal bands of Broca (VDB) and the horizontal diagonal bands of Broca (HDB) nuclei of the basal forebrain of mice 8 weeks after injection with UII-SAP or Blank-SAP (*P* < 0.0001, two-way ANOVA, Bonferroni’s multiple comparisons test; MS: *P* = 0.0004, VDB: *P* = 0.0001, HDB: *P* = 0.0032). **(C)** The number of parvalbumin-positive GABAergic neurons in the basal forebrain of mice 8 weeks after injection with UII-SAP or Blank-SAP (*P* = 0.9210, Student’s unpaired t-test). \**P* < 0.05; \*\**P* < 0.01; \*\*\**P* < 0.001; n.s., non-significant. Results are presented as mean ± s.e.m. **(D)** The percentage of time spent in the novel arm of the Y maze on test, compared to the familiar (Fam) arm (*P* = 0.9493, two-way ANOVA, Tukey’s multiple comparison test; Blank-SAP: *P* =0.0019, UII-SAP: *P* = 0.0446). Both cMPT-lesioned mice and sham-lesioned wildtype mice displayed a preference for the novel arm. **(E)** The percentage of time spent examining a novel object compared to a familiar object (*P* > 0.9999, two-way ANOVA, Tukey’s multiple comparison test; Blank-SAP: *P* = 0.0173, UII-SAP: *P* = 0.0020). Both cMPT-lesioned mice and sham-lesioned mice displayed a preference for the novel object, with no difference in the time spent in each arm between the cMPT-lesioned wildtype mice and sham-lesioned mice.

**Supplementary Figure 4.**
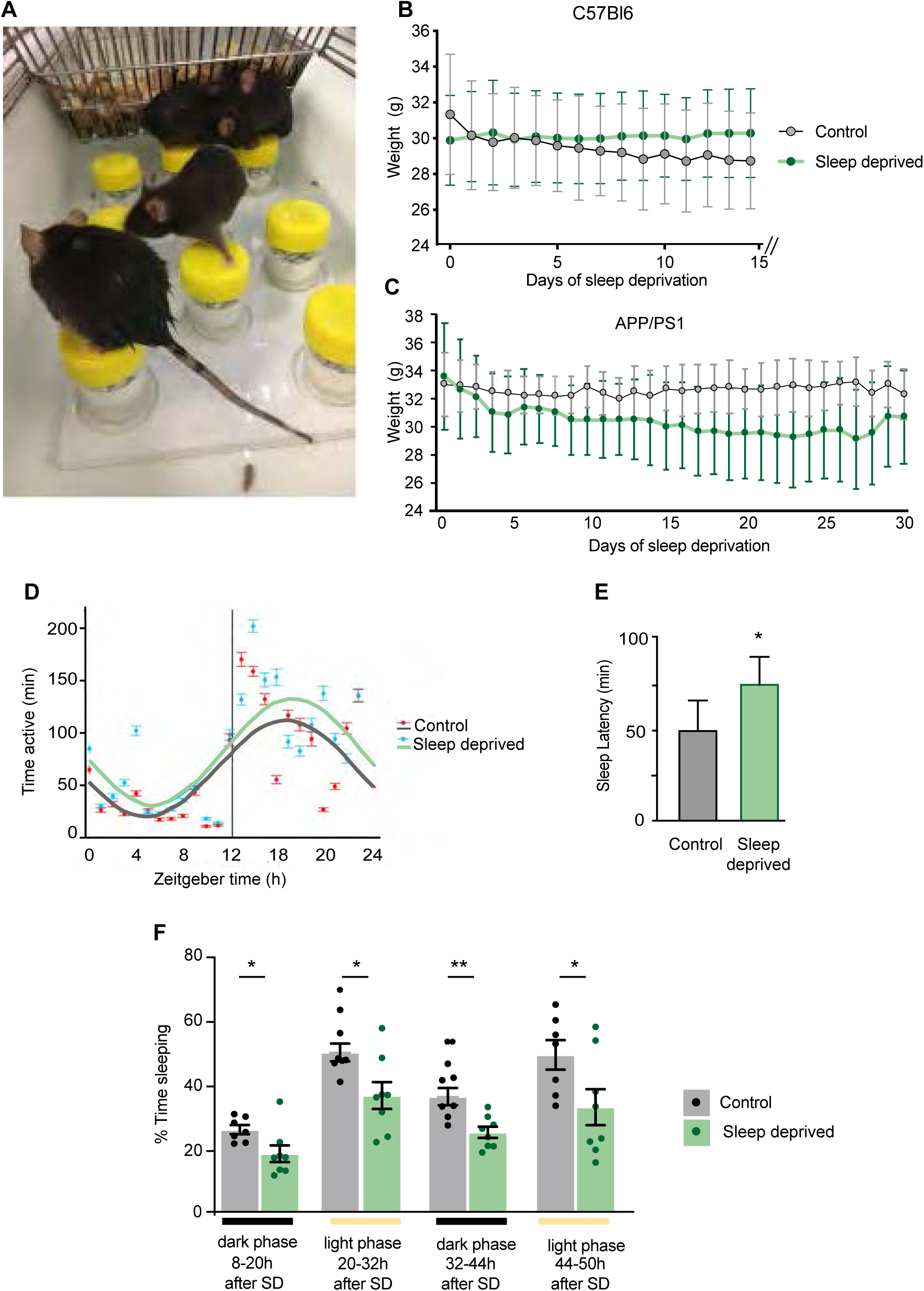
Lasting effects of the sleep deprivation paradigm. **(A)** Photograph of the sleep deprivation cages. (**B-C**) Weight loss of sleep-deprived and control wildtype **(B)** and APP/PS1 **(C)** mice. **(D)** Circadian rhythms of APP/PS1 mice in the 24 hours immediately following sleep deprivation. **(E)** Sleep latency of control and sleep-deprived mice immediately after being transferred from sleep-deprivation cages to home cages on the last sleep-deprivation day. **(F)** Sleep analysis of APP/PS1 mice in the 3 days following sleep deprivation. Sleep-deprived mice have reduced sleep time (*P* values left to right are: 0.0298, 0.0224, 0.0035, 0.0462 [and number of sleep epochs (not shown) *P* = 0.0299, 0.224, 0.0035, 0.0303]) in both light and dark periods. \**P* < 0.05 \*\**P* < 0.01; Unpaired t-tests. Results are presented as mean ± s.d in B and C and mean ± s.e.m in D and E.

**Supplementary Figure 5. Inflammation in sleep-deprived APP/PS1 mice is not increased**

Area of CD68-postive microglia in sensory cortex (**A**) and hippocampus (**B**) of control or sleep-deprived APP/PS1 mice.

Area of GFAP-positive astrocytes in sensory cortex (**C**) and hippocampus (**D**) of control or sleep-deprived APP/PS1 mice. A significant reduction in astrocytosis was found in the hippocampus of sleep-deprived mice compared to controls (* *P=*0.027, unpaired two-way t-test). Results are presented as mean ± s.e.m.

**Supplementary Figure 6.**
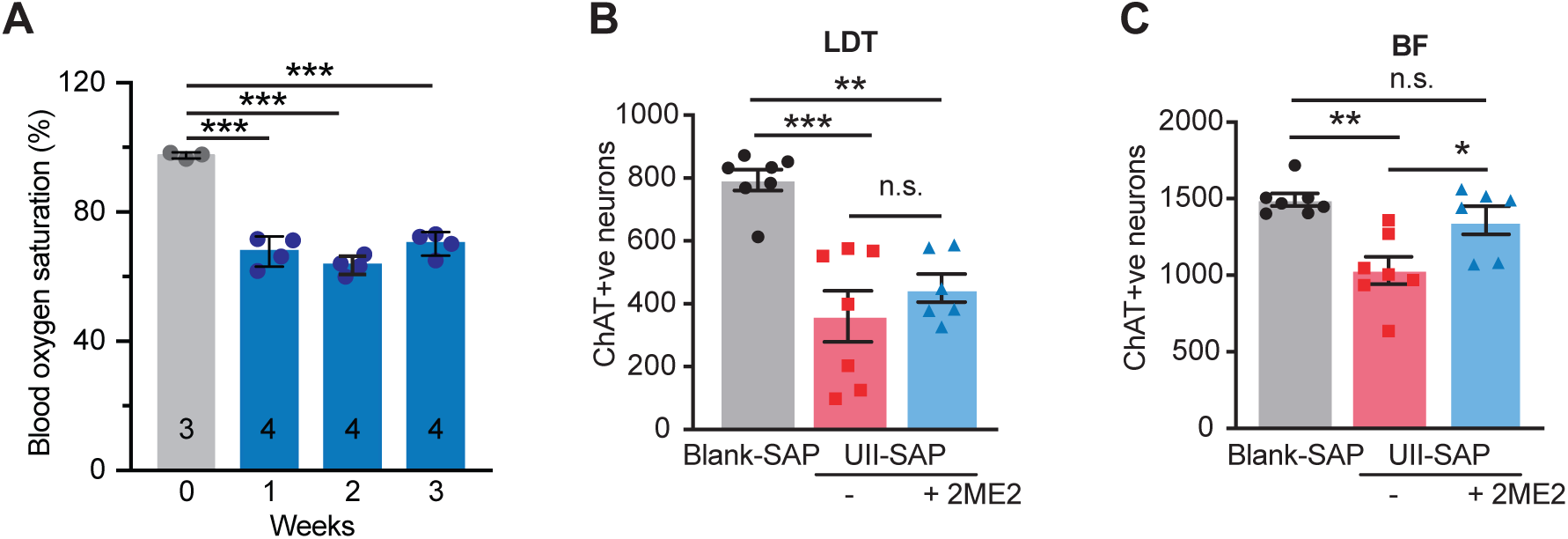
2ME2 treatment protects from OSA-induced cBF degeneration. **(A)** Blood oximetry measurements of mice during each of the 4 weeks of chronic hypoxia exposure. The number of cMPT **(B)** and cBF **(C)** neurons in a second cohort of mice injected with either Blank-SAP or UII-SAP treated with daily 15mg/kg 2ME2 or vehicle for 27 days (cBF: Blank-SAP vs. UII-SAP: *P* = 0.0011, Blank-SAP vs. 2ME2 treated: *P* = 0.020, UII-SAP vs. 2ME2 treated: *P* = 0.4549. As in Fig 9B, 2ME2 treatment protects cBF neurons from the effects of OSA. \**P* < 0.05; \*\**P* < 0.01; \*\*\**P* < 0.001; n.s., non-significant, one-way ANOVA with Tukey’s multiple comparison test. Results are presented as mean ± s.e.m.

**Supplementary Figure 7.**
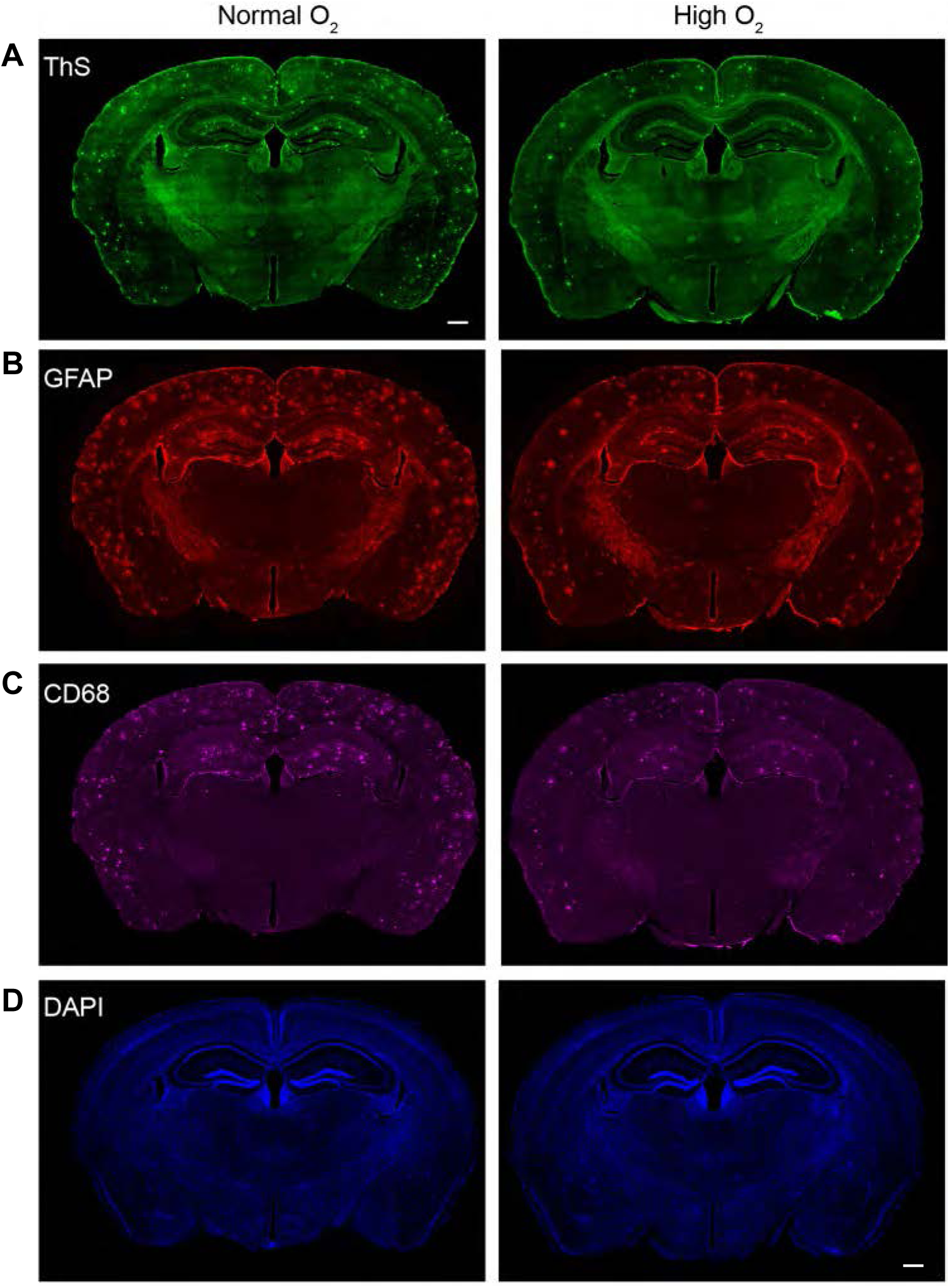
‘CPAP’ rescues APP/PS1 OSA mice from exacerbated AD pathology. Representative images of **(A)** thioflavin-S (ThS)-positive Aβ plaques, **(B)** GFAP-positive astrocytes, **(C)** CD68-positive microglia, and (**D**) DAPI-positive nuclei in coronal sections containing the neocortex and hippocampus from APP/PS1 mice placed in 40% oxygenated (high O_2_) or normoxia (normal O_2_) for 8 hours a day during the sleep period for 6 weeks, starting 2 weeks after injection with UII-SAP. Scale bar = 200 μm.

## REFERENCES

Adams N, Strauss M, Schluchter M, Redline S (2001) Relation of measures of sleep-disordered breathing to neuropsychological functioning. Am J Respir Crit Care Med 163:1626–1631.

Ancoli-Israel S, Palmer BW, Cooke JR, Corey-Bloom J, Fiorentino L, Natarajan L, Liu L, Ayalon L, He F, Loredo JS (2008) Cognitive effects of treating obstructive sleep apnoea in Alzheimer’s disease: a randomized controlled study. J Am Geriatr Soc 56:2076–2081.

Arias-Cavieres A, Khuu MA, Nwakudu CU, Barnard JE, Dalgin G, Garcia AJ (2019) A role for Hypoxia Inducible Factor 1a (HIF1a) in intermittent hypoxia-dependent changes to spatial memory and synaptic plasticity. bioRxiv:595975.

Atkins L, Bourgeat P, Fripp J, Coulson EJ (2016) Basal forebrain degeneration precedes cognitive decline in obstructive sleep apnoea patients. In: 15th National Conference of Emerging Researchers in Ageing. Canberra, Australia.

Ballinger EC, Ananth M, Talmage DA, Role LW (2016) Basal forebrain cholinergic circuits and signaling in cognition and cognitive decline. Neuron 91:1199–1218.

Barrett GL, Reid CA, Tsafoulis C, Zhu W, Williams DA, Paolini AG, Trieu J, Murphy M (2010) Enhanced spatial memory and hippocampal long-term potentiation in p75 neurotrophin receptor knockout mice. Hippocampus 20:145–152.

Baxter MG, Chiba AA (1999) Cognitive functions of the basal forebrain. Curr Opin Neurobiol 9:178–183.

Bellingham MC, Berger AJ (1996) Presynaptic depression of excitatory synaptic inputs to rat hypoglossal motoneurons by muscarinic M2 receptors. J Neurophysiol 76:3758–3770.

Boskovic Z, Alfonsi F, Rumballe BA, Fonseka S, Windels F, Coulson EJ (2014) The role of p75_NTR_ in cholinergic basal forebrain structure and function. J Neurosci 34:13033–13038.

Boskovic Z, Meier S, Wang Y-R, Milne MR, Onraet T, Tedoldi A, Coulson EJ (2019) Regulation of cholinergic basal forebrain development, connectivity, and function by neurotrophin receptors. Neuronal Signaling 3.

Boskovic Z, Milne M, Qian L, Clifton H, McGovern AE, Turnbull MT, Mazzone SB, Coulson EJ (2018) Cholinergic basal forebrain neurons regulate fear extinction consolidation through p75 neurotrophin receptor signaling. Trans Psych. 8:199-

Cavadas MAS, Cheong A, Taylor CT (2017) The regulation of transcriptional repression in hypoxia. Exp Cell Res 356:173–181.

Chiesa PA, Cavedo E, Grothe MJ, Houot M, Teipel SJ, Potier MC, Habert MO, Lista S, Dubois B, Hampel H (2019) Relationship between basal forebrain resting-state functional connectivity and brain amyloid-beta deposition in cognitively intact older adults with subjective memory complaints. Radiology 290:167–176.

Cimadevilla JM, Kaminsky Y, Fenton A, Bures J (2000) Passive and active place avoidance as a tool of spatial memory research in rats. J Neurosci Methods 102:155–164.

Clark RJ, Fischer H, Dempster L, Daly NL, Rosengren KJ, Nevin ST, Meunier FA, Adams DJ, Craik DJ (2005) Engineering stable peptide toxins by means of backbone cyclization: stabilization of the alpha-conotoxin MII. Proc Natl Acad Sci U S A 102:13767–13772.

Clark SD, Nothacker HP, Wang Z, Saito Y, Leslie FM, Civelli O (2001) The urotensin II receptor is expressed in the cholinergic mesopontine tegmentum of the rat. Brain Res 923:120–127.

Daulatzai MA (2014) Role of stress, depression, and aging in cognitive decline and Alzheimer’s disease. Curr Top Behav Neurosci 18:265–296.

Daulatzai MA (2015) Evidence of neurodegeneration in obstructive sleep apnoea: Relationship between obstructive sleep apnoea and cognitive dysfunction in the elderly. J Neurosci Res 93:1778–1794.

Dayan F, Roux D, Brahimi-Horn MC, Pouyssegur J, Mazure NM (2006) The oxygen sensor factor-inhibiting hypoxia-inducible factor-1 controls expression of distinct genes through the bifunctional transcriptional character of hypoxia-inducible factor-1alpha. Cancer Res 66:3688–3698.

Elias A, Cummins T, Tyrrell R, Lamb F, Dore V, Williams R, Rosenfeld JV, Hopwood M, Villemagne VL, Rowe CC (2018) Risk of alzheimer’s disease in obstructive sleep apnoea syndrome: amyloid-beta and tau Imaging. J Alzheimers Dis 66:733–741.

Findley LJ, Barth JT, Powers DC, Wilhoit SC, Boyd DG, Suratt PM (1986) Cognitive impairment in patients with obstructive sleep apnoea and associated hypoxemia. Chest 90:686–690.

Fisher A, Bezprozvanny I, Wu L, Ryskamp DA, Bar-Ner N, Natan N, Brandeis R, Elkon H, Nahum V, Gershonov E, LaFerla FM, Medeiros R (2016) AF710B, a novel M1/sigma1 agonist with therapeutic efficacy in animal models of Alzheimer’s Disease. Neurodegener Dis 16:95–110.

Gil-Bea FJ, Gerenu G, Aisa B, Kirazov LP, Schliebs R, Ramirez MJ (2012) Cholinergic denervation exacerbates amyloid pathology and induces hippocampal atrophy in Tg2576 mice. Neurobiol Dis 48:439–446.

Greferath U, Bennie A, Kourakis A, Barrett GL (2000) Impaired spatial learning in aged rats is associated with loss of p75-positive neurons in the basal forebrain. Neuroscience 100:363–373.

Grothe M, Heinsen H, Teipel S (2013) Longitudinal measures of cholinergic forebrain atrophy in the transition from healthy aging to Alzheimer’s disease. Neurobiol Aging 34:1210–1220.

Grothe MJ, Ewers M, Krause B, Heinsen H, Teipel SJ, Alzheimer’s Disease Neuroimaging I (2014a) Basal forebrain atrophy and cortical amyloid deposition in nondemented elderly subjects. Alzheimers Dement 10:S344–353.

Grothe MJ, Schuster C, Bauer F, Heinsen H, Prudlo J, Teipel SJ (2014b) Atrophy of the cholinergic basal forebrain in dementia with Lewy bodies and Alzheimer’s disease dementia. J Neurol 261:1939–1948.

Hamlin AS, Windels F, Boskovic Z, Sah P, Coulson EJ (2013) Lesions of the basal forebrain cholinergic system in mice disrupt idiothetic navigation. PLoS One 8:e53472.

Hampel H, Mesulam MM, Cuello AC, Khachaturian AS, Vergallo A, Farlow MR, Snyder PJ, Giacobini E, Khachaturian ZS (2019) Revisiting the cholinergic hypothesis in Alzheimer’s disease: Emerging evidence from translational and clinical research. J Prevention Alz dis 6:2–15.

Hardy J, Selkoe DJ (2002) The amyloid hypothesis of Alzheimer’s disease: progress and problems on the road to therapeutics. Science 297:353–356.

Hartig W, Saul A, Kacza J, Grosche J, Goldhammer S, Michalski D, Wirths O (2014) Immunolesion-induced loss of cholinergic projection neurones promotes beta-amyloidosis and tau hyperphosphorylation in the hippocampus of triple-transgenic mice. Neuropathol Appl Neurobiol 40:106–120.

Hernandez AB, Kirkness JP, Smith PL, Schneider H, Polotsky M, Richardson RA, Hernandez WC, Schwartz AR (2012) Novel whole body plethysmography system for the continuous characterization of sleep and breathing in a mouse. Journal of applied physiology (Bethesda, Md : 1985) 112:671–680.

Hort J, Laczo J, Vyhnalek M, Bojar M, Bures J, Vlcek K (2007) Spatial navigation deficit in amnestic mild cognitive impairment. Proc Natl Acad Sci U S A 104:4042–4047.

Hort J, Andel R, Mokrisova I, Gazova I, Amlerova J, Valis M, Coulson EJ, Harrison J, Windisch M, Laczo J (2014) Effect of Donepezil in Alzheimer disease can be measured by a computerized human analog of the Morris Water Maze. Neurodegen Dis 13:192–196.

Ibanez CF, Simi A (2012) p75 neurotrophin receptor signaling in nervous system injury and degeneration: paradox and opportunity. Trends Neurosci 35:431–440.

Incalzi RA, Marra C, Salvigni BL, Petrone A, Gemma A, Selvaggio D, Mormile F (2004) Does cognitive dysfunction conform to a distinctive pattern in obstructive sleep apnoea syndrome? J Sleep Res 13:79–86.

Irmak SO, de Lecea L (2014) Basal forebrain cholinergic modulation of sleep transitions. Sleep 37:1941–1951.

Iyalomhe O, Swierczek S, Enwerem N, Chen Y, Adedeji MO, Allard J, Ntekim O, Johnson S, Hughes K, Kurian P, Obisesan TO (2017) The role of Hypoxia-Inducible Factor 1 in mild cognitive impairment. Cell Mol Neurobiol 37:969–977.

Jankowsky JL, Fadale DJ, Anderson J, Xu GM, Gonzales V, Jenkins NA, Copeland NG, Lee MK, Younkin LH, Wagner SL, Younkin SG, Borchelt DR (2004) Mutant presenilins specifically elevate the levels of the 42 residue beta-amyloid peptide in vivo: evidence for augmentation of a 42-specific gamma secretase. Hum Mol Genet 13:159–170.

Jegou S, Cartier D, Dubessy C, Gonzalez BJ, Chatenet D, Tostivint H, Scalbert E, LePrince J, Vaudry H, Lihrmann I (2006) Localization of the urotensin II receptor in the rat central nervous system. J Comp Neurol 495:21–36.

Kerbler GM, Fripp J, Rowe CC, Villemagne VL, Salvado O, Rose S, Coulson EJ, Alzheimer’s Disease Neuroimaging I (2015a) Basal forebrain atrophy correlates with amyloid beta burden in Alzheimer’s disease. NeuroImage Clinical 7:105–113.

Kerbler GM, Nedelska Z, Fripp J, Laczo J, Vyhnalek M, Lisy Jvri, Hamlin AS, Rose S, Hort J, Coulson EJ (2015b) Basal forebrain atrophy contributes to allocentric navigation impairment in Alzheimer’s disease patients. Front Aging Neurosci 7.

Knowles JK, Rajadas J, Nguyen TV, Yang T, LeMieux MC, Vander Griend L, Ishikawa C, Massa SM, Wyss-Coray T, Longo FM (2009) The p75 neurotrophin receptor promotes amyloid-beta1-42-induced neuritic dystrophy in vitro and in vivo. J Neurosci 29:10627–10637.

Laursen B, Mork A, Plath N, Kristiansen U, Bastlund JF (2014) Impaired hippocampal acetylcholine release parallels spatial memory deficits in Tg2576 mice subjected to basal forebrain cholinergic degeneration. Brain Res 1543:253–262.

Le Moan N, Houslay DM, Christian F, Houslay MD, Akassoglou K (2011) Oxygen-dependent cleavage of the p75 neurotrophin receptor triggers stabilization of HIF-1alpha. Mol Cell 44:476–490.

Livingston G et al. (2017) Dementia prevention, intervention, and care. Lancet 390:2673–2734.

Longo FM, Massa SM (2013) Small-molecule modulation of neurotrophin receptors: a strategy for the treatment of neurological disease. Nature reviews Drug discovery 12:507–525.

Mander BA, Winer JR, Jagust WJ, Walker MP (2016) Sleep: A Novel Mechanistic Pathway, Biomarker, and Treatment Target in the Pathology of Alzheimer’s Disease? Trends Neurosci 39:552–566.

Martinez CA, Kerr B, Jin C, Cistulli PA, Cook KM (2019) Obstructive sleep apnoea activates HIF-1 in a hypoxia dose-dependent manner in HCT116 colorectal carcinoma cells. Int J Mol Sci 20.

Mizuno S, Kameda A, Inagaki T, Horiguchi J (2004) Effects of donepezil on Alzheimer’s disease: the relationship between cognitive function and rapid eye movement sleep. Psychiatry Clin Neurosci 58:660–665.

Navarrete-Opazo A, Mitchell GS (2014) Therapeutic potential of intermittent hypoxia: a matter of dose. Am J Physiol Regul Integr Comp Physiol 307:R1181–1197.

Osorio RS, Pirraglia E, Gumb T, Mantua J, Ayappa I, Williams S, Mosconi L, Glodzik L, de Leon MJ (2014a) Imaging and cerebrospinal fluid biomarkers in the search for Alzheimer’s disease mechanisms. Neurodegener Dis 13:163–165.

Osorio RS, Pirraglia E, Aguera-Ortiz LF, During EH, Sacks H, Ayappa I, Walsleben J, Mooney A, Hussain A, Glodzik L, Frangione B, Martinez-Martin P, de Leon MJ (2011) Greater risk of Alzheimer’s disease in older adults with insomnia. J Am Geriatr Soc 59:559–562.

Osorio RS, Ayappa I, Mantua J, Gumb T, Varga A, Mooney AM, Burschtin OE, Taxin Z, During E, Spector N, Biagioni M, Pirraglia E, Lau H, Zetterberg H, Blennow K, Lu SE, Mosconi L, Glodzik L, Rapoport DM, de Leon MJ (2014b) Interaction between sleep-disordered breathing and apolipoprotein E genotype on cerebrospinal fluid biomarkers for Alzheimer’s disease in cognitively normal elderly individuals. Neurobiol Aging 35:1318–1324.

Owen JE, BenediktsdOttir B, Gislason T, Robinson SR (2019) Neuropathological investigation of cell layer thickness and myelination in the hippocampus of people with obstructive sleep apnoea. Sleep 42.

Punjabi NM (2008) The epidemiology of adult obstructive sleep apnoea. Proceedings of the American Thoracic Society 5:136–143.

Qian L, Milne MR, Shepheard S, Rogers ML, Medeiros R, Coulson EJ (2018) Removal of p75 Neurotrophin Receptor Expression from Cholinergic Basal Forebrain Neurons Reduces Amyloid-beta Plaque Deposition and Cognitive Impairment in Aged APP/PS1 Mice. Mol Neurobiol.

Ramirez JM, Garcia AJ, 3rd, Anderson TM, Koschnitzky JE, Peng YJ, Kumar GK, Prabhakar NR (2013) Central and peripheral factors contributing to obstructive sleep apnoeas. Respir Physiol Neurobiol 189:344–353.

Ramos-Rodriguez JJ, Pacheco-Herrero M, Thyssen D, Murillo-Carretero MI, Berrocoso E, Spires-Jones TL, Bacskai BJ, Garcia-Alloza M (2013) Rapid beta-amyloid deposition and cognitive impairment after cholinergic denervation in APP/PS1 mice. J Neuropathol Exp Neurol 72:272–285.

Richards KC, Gooneratne N, Dicicco B, Hanlon A, Moelter S, Onen F, Wang Y, Sawyer A, Weaver T, Lozano A, Carter P, Johnson J (2019) CPAP adherence may slow 1-Year cognitive decline in older adults with mild cognitive impairment and apnoea. J Am Geriatr Soc 67:558–564.

Rosenzweig I, Williams SC, Morrell MJ (2014) The impact of sleep and hypoxia on the brain: potential mechanisms for the effects of obstructive sleep apnoea. Curr Opin Pulm Med 20:565–571.

Rossner S, Ueberham U, Schliebs R, Perez-Polo JR, Bigl V (1998) The regulation of amyloid precursor protein metabolism by cholinergic mechanisms and neurotrophin receptor signaling. Prog Neurobiol 56:541–569.

Row BW, Kheirandish L, Cheng Y, Rowell PP, Gozal D (2007) Impaired spatial working memory and altered choline acetyltransferase (CHAT) immunoreactivity and nicotinic receptor binding in rats exposed to intermittent hypoxia during sleep. Behav Brain Res 177:308–314.

Sanchez-Ortiz E, Yui D, Song D, Li Y, Rubenstein JL, Reichardt LF, Parada LF (2012) TrkA gene ablation in basal forebrain results in dysfunction of the cholinergic circuitry. J Neurosci 32:4065–4079.

Schmitz TW, Nathan Spreng R, Alzheimer’s Disease Neuroimaging I (2016) Basal forebrain degeneration precedes and predicts the cortical spread of Alzheimer’s pathology. Nat Commun 7:13249.

Shiota S, Takekawa H, Matsumoto SE, Takeda K, Nurwidya F, Yoshioka Y, Takahashi F, Hattori N, Tabira T, Mochizuki H, Takahashi K (2013) Chronic intermittent hypoxia/reoxygenation facilitate amyloid-beta generation in mice. J Alzheimers Dis 37:325–333.

Sotthibundhu A, Sykes AM, Fox B, Underwood CK, Thangnipon W, Coulson EJ (2008) Beta-amyloid1-42 induces neuronal death through the p75 neurotrophin receptor. J Neurosci 28:3941–3946.

Spira AP, Gottesman RF (2017) Sleep disturbance: an emerging opportunity for Alzheimer’s disease prevention? Int Psychogeriatr 29:529–531.

Spira AP, Yager C, Brandt J, Smith GS, Zhou Y, Mathur A, Kumar A, Brasic JR, Wong DF, Wu MN (2014) Objectively measured sleep and beta-amyloid burden in older adults: A pilot Study. SAGE Open Medicine 2.

Tarasoff-Conway JM, Carare RO, Osorio RS, Glodzik L, Butler T, Fieremans E, Axel L, Rusinek H, Nicholson C, Zlokovic BV, Frangione B, Blennow K, Menard J, Zetterberg H, Wisniewski T, de Leon MJ (2015) Clearance systems in the brain-implications for Alzheimer disease. Nat Rev Neurol 11:457–470.

Teipel S, Heinsen H, Amaro E, Jr., Grinberg LT, Krause B, Grothe M, Alzheimer’s Disease Neuroimaging I (2014) Cholinergic basal forebrain atrophy predicts amyloid burden in Alzheimer’s disease. Neurobiol Aging 35:482–491.

Tong B, Pantazopoulou V, Johansson E, Pietras A (2018) The p75 neurotrophin receptor enhances HIF-dependent signaling in glioma. Exp Cell Res 371:122–129.

Turnbull MT, Boskovic Z, Coulson EJ (2018) Acute down-regulation of BDNF signaling does not replicate exacerbated amyloid-beta levels and cognitive impairment induced by cholinergic basal forebrain lesion. Front Mol Neurosci 11:51.

Van Dort CJ, Zachs DP, Kenny JD, Zheng S, Goldblum RR, Gelwan NA, Ramos DM, Nolan MA, Wang K, Weng FJ, Lin Y, Wilson MA, Brown EN (2015) Optogenetic activation of cholinergic neurons in the PPT or LDT induces REM sleep. Proc Natl Acad Sci U S A 112:584–589.

Verstraeten E, Cluydts R (2004) Executive control of attention in sleep apnoea patients: theoretical concepts and methodological considerations. Sleep Med Rev 8:257–267.

Villa JC, Chiu D, Brandes AH, Escorcia FE, Villa CH, Maguire WF, Hu CJ, de Stanchina E, Simon MC, Sisodia SS, Scheinberg DA, Li YM (2014) Nontranscriptional role of Hif-1alpha in activation of gamma-secretase and notch signaling in breast cancer. Cell Rep 8:1077–1092.

Wang G, Grone B, Colas D, Appelbaum L, Mourrain P (2011) Synaptic plasticity in sleep: learning, homeostasis and disease. Trends Neurosci 34:452–463.

Ward NR, Roldao V, Cowie MR, Rosen SD, McDonagh TA, Simonds AK, Morrell MJ (2013) The effect of respiratory scoring on the diagnosis and classification of sleep disordered breathing in chronic heart failure. Sleep 36:1341–1348.

Welt T, Kulic L, Hoey SE, McAfoose J, Spani C, Chadha AS, Fisher A, Nitsch RM (2015) Acute Effects of muscarinic M1 receptor modulation on AbetaPP metabolism and amyloid-beta levels in vivo: A Microdialysis Study. J Alzheimers Dis 46:971–982.

Wesierska M, Dockery C, Fenton AA (2005) Beyond memory, navigation, and inhibition: behavioral evidence for hippocampus-dependent cognitive coordination in the rat. J Neurosci 25:2413–2419.

Wollam ME, Weinstein AM, Saxton JA, Morrow L, Snitz B, Fowler NR, Suever Erickson BL, Roecklein KA, Erickson KI (2015) Genetic risk score predicts late-life cognitive impairment. Journal of Aging Research 2015:267062.

Woolf NJ, Butcher LL (1986) Cholinergic systems in the rat brain: III. Projections from the pontomesencephalic tegmentum to the thalamus, tectum, basal ganglia, and basal forebrain. Brain Res Bull 16:603–637.

Xiao C, Cho JR, Zhou C, Treweek JB, Chan K, McKinney SL, Yang B, Gradinaru V (2016) Cholinergic mesopontine signals govern locomotion and reward through dissociable midbrain pathways. Neuron 90:333–347.

Yaffe K, Falvey CM, Hoang T (2014) Connections between sleep and cognition in older adults. Lancet Neurol 13:1017–1028.

Yaffe K, Laffan AM, Harrison SL, Redline S, Spira AP, Ensrud KE, Ancoli-Israel S, Stone KL (2011) Sleep-disordered breathing, hypoxia, and risk of mild cognitive impairment and dementia in older women. JAMA 306:613–619.

Yun CH, Lee HY, Lee SK, Kim H, Seo HS, Bang SA, Kim SE, Greve DN, Au R, Shin C, Thomas RJ (2017) Amyloid Burden in Obstructive Sleep Apnoea. J Alzheimers Dis 59:21–29.

Zhang B, Veasey SC, Wood MA, Leng LZ, Kaminski C, Leight S, Abel T, Lee VM, Trojanowski JQ (2005) Impaired rapid eye movement sleep in the Tg2576 APP murine model of Alzheimer’s disease with injury to pedunculopontine cholinergic neurons. The American Journal of Pathology 167:1361–1369.

Zhang X, Zhou K, Wang R, Cui J, Lipton SA, Liao FF, Xu H, Zhang YW (2007) Hypoxia-inducible factor 1alpha (HIF-1alpha)-mediated hypoxia increases BACE1 expression and beta-amyloid generation. J Biol Chem 282:10873–10880.

Zhao Z, Zhao X, Veasey SC (2017) Neural consequences of chronic short sleep: Reversible or lasting? Front Neurol 8:235.

Zhu X, Zhao Y (2018) Sleep-disordered breathing and the risk of cognitive decline: a meta-analysis of 19,940 participants. Sleep & Breathing; Schlaf & Atmung 22:165–173.

Zuroff L, Daley D, Black KL, Koronyo-Hamaoui M (2017) Clearance of cerebral Abeta in Alzheimer’s disease: reassessing the role of microglia and monocytes. Cell Mol Life Sci 74:2167–2201.

